# Self-organization and information transfer in Antarctic krill swarms

**DOI:** 10.1101/2021.01.19.427357

**Authors:** Alicia L. Burns, Timothy M. Schaerf, Joseph T. Lizier, So Kawaguchi, Martin Cox, Rob King, Jens Krause, Ashley J.W. Ward

**Affiliations:** School of Life and Environmental Sciences, University of Sydney, Sydney, NSW, Australia; School of Science and Technology, University of New England, Armidale, NSW, Australia; Complex Systems Research Group, Faculty of Engineering and IT, University of Sydney, Sydney, NSW, Australia; Australian Antarctic Division, Kingston, Tasmania, Australia; Department of Biology and Ecology of Fishes, Leibniz-Institute of Freshwater Ecology and Inland Fisheries, Müggelseedamm 310, Berlin 12587, Germany; Faculty of Life Science, Humboldt-Universität zu Berlin, Invalidenstrasse 42, Berlin 10115, Germany; Taronga Institute of Science and Learning, Taronga Conservation Society Australia, Mosman, NSW, Australia

## Abstract

Antarctic krill swarms are one of the largest known animal aggregations. However, despite being the keystone species of the Southern Ocean, little is known about how swarms are formed and maintained, and we lack a detailed understanding of the local interactions between individuals that provide the basis for these swarms. Here we analyzed the trajectories of captive, wild-caught krill in 3D to determine individual level interaction rules and quantify patterns of information flow. Our results suggest krill operate a novel form of collective organization, with measures of information flow and individual movement adjustments expressed most strongly in the vertical dimension, a finding not seen in other swarming species. In addition, local directional alignment with near neighbors, and strong regulation of both direction and speed relative to the positions of groupmates suggest social factors are vital to the formation and maintenance of swarms. This research represents a first step in understanding the fundamentally important swarming behavior of krill.

---

The Antarctic krill (*Euphausia superba*) is one of the most abundant and important animal species, and is often described as the keystone species of the Southern Ocean Ecosystem. The aggregation of krill into swarms is thought to be a major part of their success, providing safety in numbers (*1*), the ability to track nutrient gradients, and an increase in swimming efficiency, leading to vital energy savings (*2*). Given the crucial importance of swarming to the survival of krill, a number of studies have examined the structure and function of krill groups in both the laboratory and the field (*3,4*). Nonetheless, we currently lack a detailed understanding of how these swarms are formed and maintained.

Previous work on grouping animals, mostly studied in 2-dimensions, has shown that group-level patterns of collective motion emerge through the repeated interactions that occur between individual animals within the group. These interactions can often be distilled to a simple set of heuristics, including mechanisms for one or more of the following: close range repulsion, long range attraction, and localized alignment. Together, such interactions are known as rules of interaction, or rules of motion. These rules generally describe how individuals adjust their behavior based on the relative locations and actions of groupmates, providing insight to the global structuring of animal collectives. Similarly, information transfer arises in interactions between near neighbors relating to changes in speed or heading direction, resulting in individuals sequentially adapting their trajectories in time. Efficient information transfer among group members is critical to the effectiveness of collective behavior. Information-theoretic measures, in particular transfer entropy, are now increasingly being employed to quantify information transfer in biological systems (*5-7*).

Using tracking data collected from free-swimming, captive Antarctic krill, we provide the first analysis of the interactions and information flow between individuals in three dimensions. Describing the rules of motion employed by krill in swarms, and mapping patterns of information transfer, represents an essential first step in developing a quantitative understanding of swarming in this critically important species. We determined the average rules of interaction used by individual krill to adjust their speed and heading as a function of the relative location and speed of near neighbors, adapting methods first fully developed in Herbert-Read et al.(*8*) and Katz et al.(*9*) to the study of three-dimensional collective movement.

## Results

There was a high probability of observing neighbors in close proximity to a focal individual (Fig.1, left column), within an area of local alignment extending approximately 100 - 200 mm (between 2.5 and 5 krill body lengths) from the focal to its near neighbors (Fig.1, right column). The peak occurrence of near neighbors occurs alongside the focal individual on the horizontal plane relative to both its direction of motion and the component of gravity perpendicular to the direction of motion.

Focal individuals adjusted their speed in relation to the position of near neighbors, accelerating when near neighbors were in front, or behind them, and slowing when near neighbors were above or below (Fig 2). Krill tended to swim more rapidly when their neighbors were positioned above them (Fig.2, left column).

**Fig 1:**
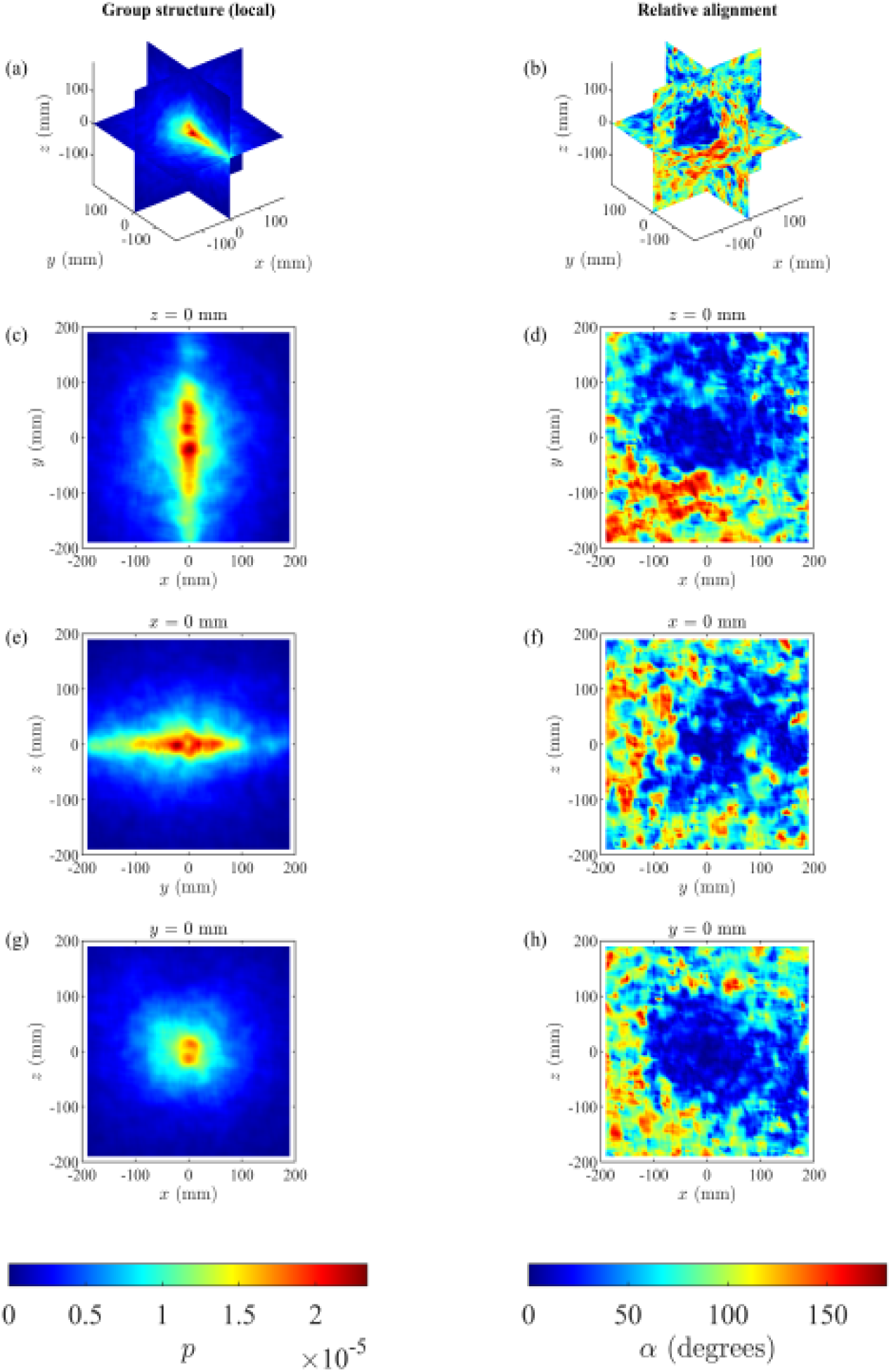
The statistical density of neighboring krill relative to a focal individual (left column), and mean angular difference in travelling direction, *α* in degrees (right column), of neighboring krill relative to a focal individual. In all panels, the focal individual is positioned at (0, 0, 0) travelling parallel to the positive *x*-axis, with the component of gravity perpendicular to the focal individual aligned with the negative *z*-axis. (a) and (b) show central slices through a cubic volume where *x* = 0, *y* = 0, and *z* = 0 (mm); (c) and (d), top down view, with the focal travelling left to right; (e) and (f) front view, with the focal travelling out of the page; and (g) and (h), right hand side, with the focal travelling left to right. The left column shows peak occurrence of near neighbors to the left and right alongside the focal individual on the same horizontal plane. The right-hand column shows low angular differences out to front and left, as well as approximately 2-3 body lengths (BL) behind. Additionally, low angular differences in travelling direction with neighbors out to approximately 1-2 BL above and below.

**Fig 2:**
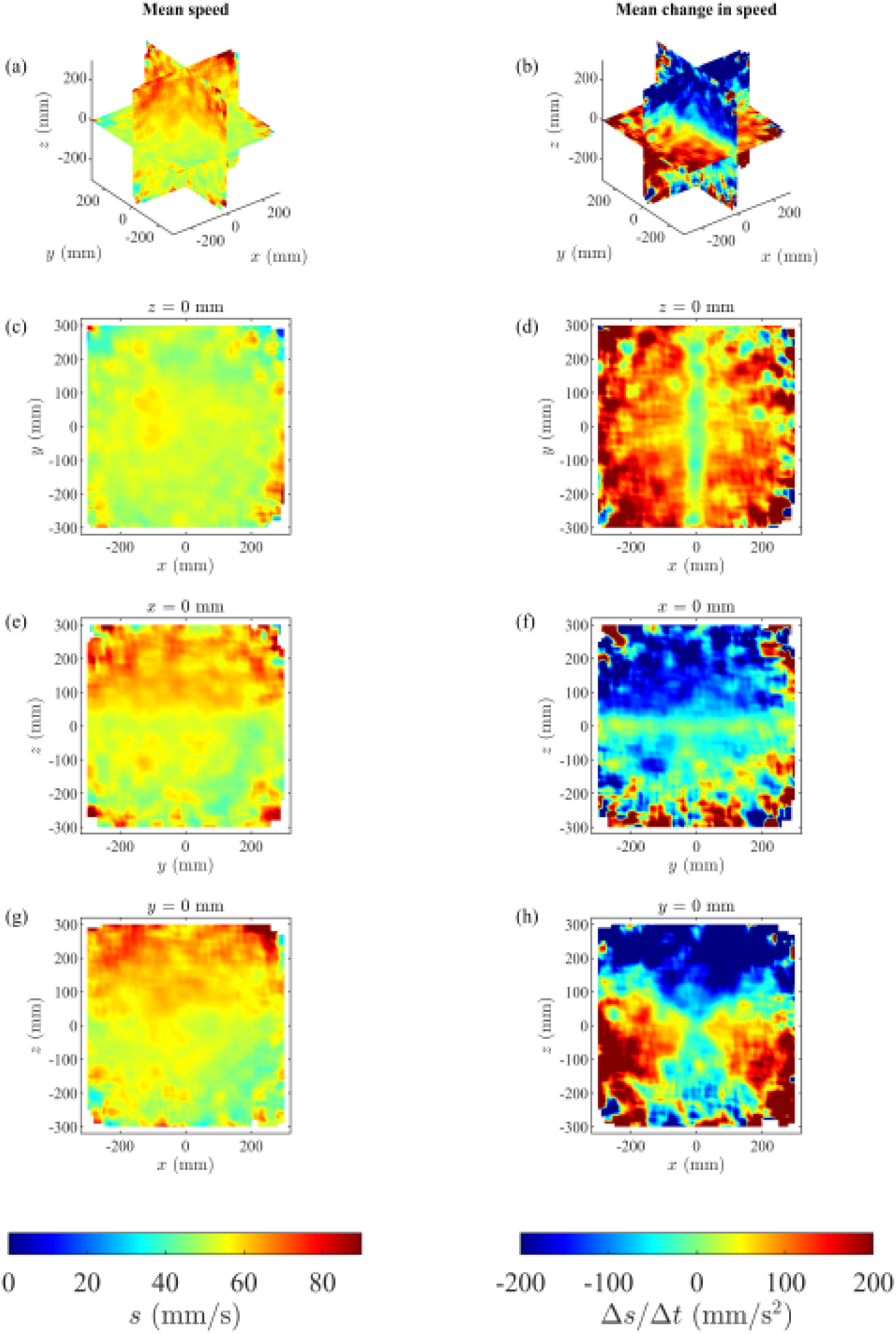
The mean speed, *s*, (left column), and change in speed, Δ*s*/Δ*t*, (right column) of krill as a function of the relative (*x, y, z*) coordinates of neighbors. Panel structure is as in Figure 1. The krill tended to travel at greater speeds on average when their neighbors occupied coordinates where *z* ≥ 50 mm (redder regions in panels (a), (e), and (g). Individual krill tended to reduce their speed when neighbors occupied the region above, where *z* ≥ 50 mm (dark blue regions in panels (b), (f), and (h)), and a smaller region below the focal individual (blue and green triangular region in panel (h)). Outside these regions, when partners occupied regions to the front and rear, and level with, or below the focal individual, then the focal individual tended to increase its speed (redder regions in panels (b), (d), and (h)).

Focal individuals also adapted their heading direction according to the position of near neighbors in both the horizontal (Fig 3, left column) and the vertical (Fig 3, right column) plane, relative to the plane of movement. Notably, when near neighbors were below and ahead, the focal turned towards them, while when near neighbors were above and in front, the focal turns upward but away from them. Broadly, focal krill exhibit little change in direction with respect to near neighbors on the same horizontal plane and within a radius of 2-3 body lengths.

**Fig 3:**
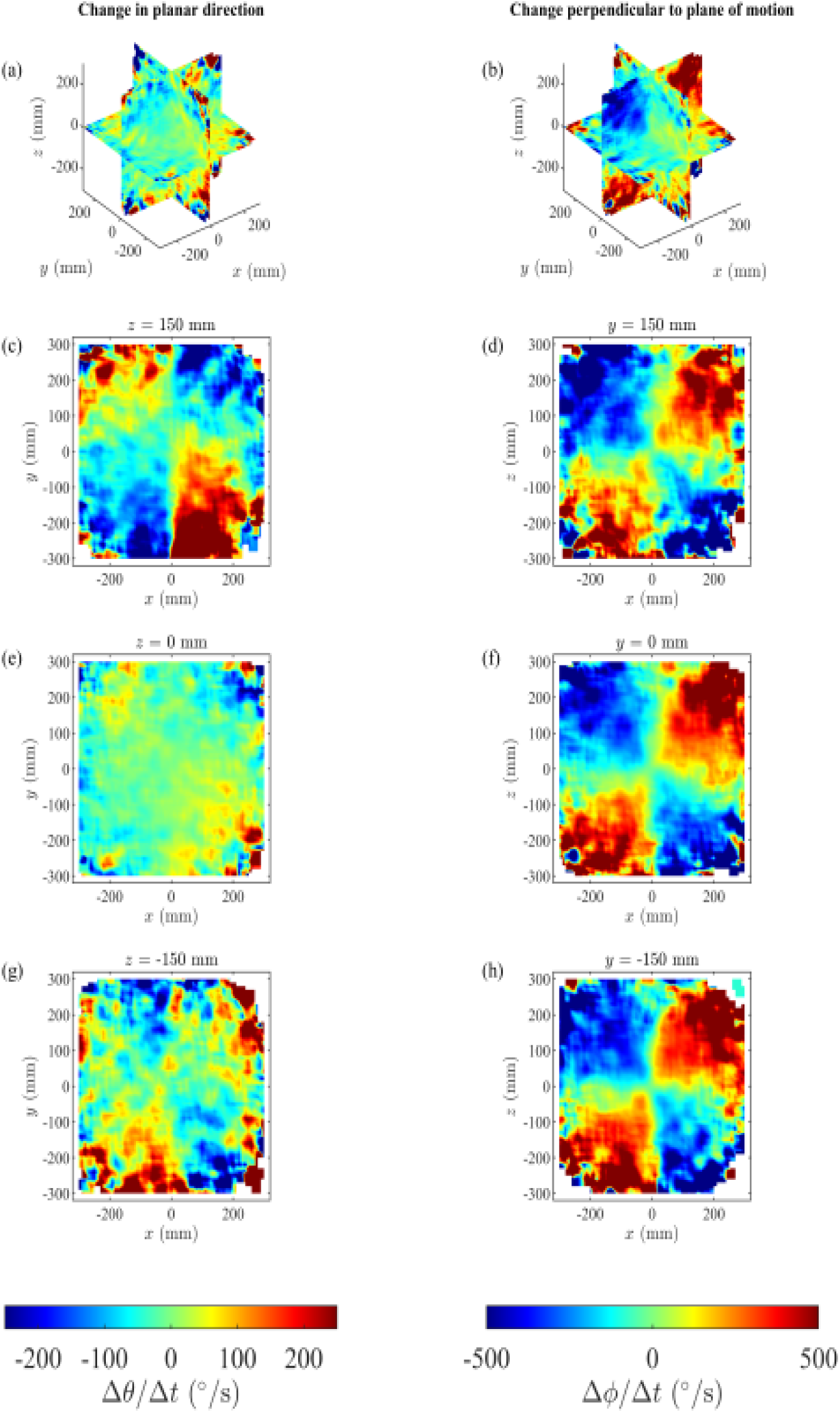
The angular components associated with the mean change in direction of krill as a function of the relative (*x, y, z*) coordinates of neighbors. Δθ/Δ*t* represents the component of turning in the plane of motion of the focal individual where *z* = 0 (left column). Positive Δθ/Δ*t* (redder regions) corresponds to leftward/anticlockwise turns, whereas negative Δθ/Δ*t* (bluer regions) corresponds to rightward/clockwise turns. (a) Δθ/Δ*t* in the planes where *x* = 0, *y* = 0, *z* = 0 (mm); (c) Δθ/Δ*t* in the plane where *z* = 150 mm (above the focal individual); (e) Δθ/Δ*t* in the plane *z* = 0 mm; (g) Δθ/Δ*t* in the plane where *z* = -150 mm (below the focal individual). Individual krill tended to make turns with rightward components when their partners were above and to their front left, or when their partners were above and to their rear right, (c). The krill tended to make turns with leftward components when their partners were above and to their front right or rear left, (c). The pattern of leftward and rightward turns reversed when partners were below (panel (g), with *z* = -150 mm). Krill would enact turns with rightward components when partners were below and to the front right or rear left, or with leftward components when partners were below and to the front left or rear right. Δ □ /Δ*t* represents the component of turning perpendicular to the plane of motion of the focal individual where *z* = 0 (right column). Positive Δ □/Δ*t* (redder regions) corresponds to upward turns, whereas negative Δ □/Δ*t* (bluer regions) corresponds to downward turns. (b) Δ □/Δt in the planes where *x* = 0, *y* = 0, and *z* = 0 (mm); (d) Δ □ □/Δ*t* in the plane where *y* = 150 mm (to the left of the focal individual); (f) Δ □/Δ*t* in the plane where *y* = 0 mm; (h) Δ □/Δ*t* in the plane where *y* = -150 mm (to the right of the focal individual). The krill tended to make turns with upward components when their neighbors were above and to the front, or below and to the rear, and turns with downward components when their partners were below and to the front or above and to the rear, irrespective of the relative left to right positions of neighbors.

Information flow, inferred from measurements of mean pairwise transfer entropy (TE), differed according to whether it was calculated from changes in heading direction, or changes in speed (see Fig 4). For both measures, information flow could be observed between focal individuals and near neighbors at all positions in the horizontal plane of movement. However, in the vertical plane of movement, information flow based on heading direction was strongest from near neighbors above and below the focal, while information flow based on speed was strongest between the focal and those near neighbors positioned in front of the focal. In general, values of transfer entropy computed on changes in heading direction were greater than that computed on changes in speed (mean Speed TE = -0.001, mean Heading TE = 0.061 nats). This, in addition to differences in the inferred interaction lag between source and target individuals for optimal TE (Speed TE: lag =3, Heading TE: lag=1), suggests a higher responsiveness and faster reactions to changes in heading direction from individuals above or below the focal, than changes in speed by those in front. For both TE computed on changes in heading direction and TE computed on changes in speed, information flow was statistically significant with reference to a null hypothesis of no directed interaction (p (surrogate > measured) < 0.01 from 100 surrogates for both Speed TE and Heading TE).

**Fig 4:**
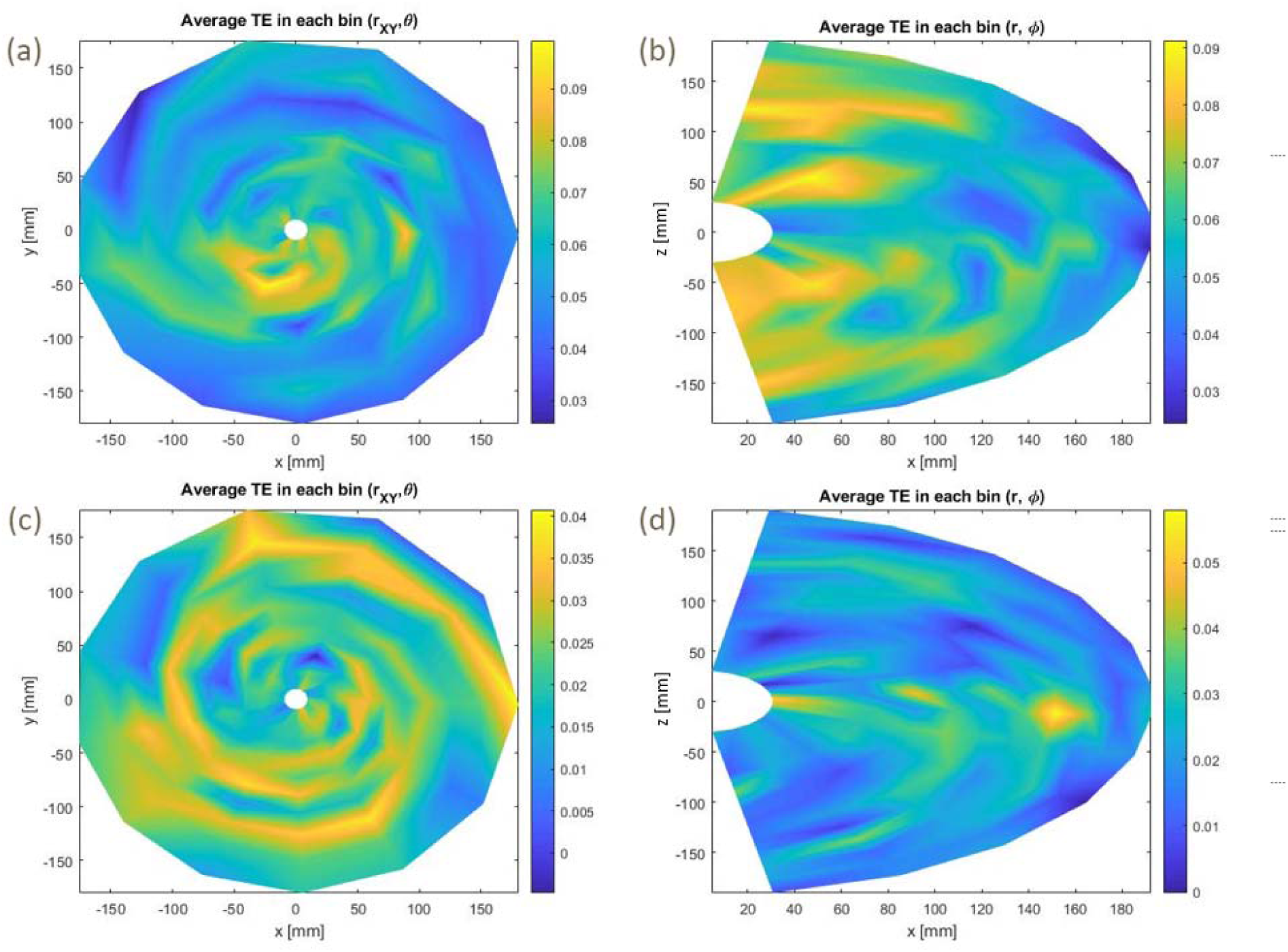
Mean pairwise transfer entropy calculated between a focal individual positioned at (*0,0*) and travelling in a direction from negative to positive *x*, and a near neighbor. Transfer entropy calculated based on changes in heading direction is shown in (a) the horizontal and (b) the vertical plane, showing localized transfer entropy from near neighbors in the same horizontal plane and peak transfer entropy from near neighbors above and below in the vertical plane. Transfer entropy calculated on changes in speed is shown in (c) the horizontal and (d) the vertical plane, showing localized transfer entropy from near neighbors in the same horizontal plane and peak transfer entropy from near neighbors lying ahead of the focal in the vertical plane. Note differences in the scale for heading versus speed transfer entropy.

## Discussion

Like other swarming species, Antarctic krill respond strongly to near neighbors and employ clear interaction rules consistent with species that demonstrate social attraction (*10*). The presence of information flow, concurrent with individual adjustments to velocity based on relative positions of neighbors, suggests that krill aggregations do not occur purely due to aggregation around food or advection by ocean currents. When near neighbors are positioned at a radius of two to three body lengths to the focal, the focal aligns with them to a large extent, which is consistent with similar observations made on shoals of fish or bird flocks moving in two dimensions (*8,11,12*). Although focal krill tend to show little change in direction when near neighbors are travelling on the same horizontal plane, they do accelerate when those near neighbors are positioned ahead or behind. Patterns of information transfer in the horizontal plane of movement, suggest focal krill are regulating speed, and particularly heading direction, potentially as a means to align with near neighbors in this plane. However, while there are some similarities between the interaction rules employed by krill and those employed by other group-living animals, there are several features that have so far not been documented, and indeed appear to be unique to krill.

Interestingly, their other responses to near neighbors, especially to those at a different vertical stratum, are qualitatively different to those reported in studies of the collective behavior of other species, including those examining the movements of animals in three dimensions, such as midges and starlings (*13,14*). Focal krill turned towards near neighbors who were ahead and below and up but away from those ahead and above. Accordingly, patterns of information transfer relating to heading direction in this vertical plane show strong interactions between focals and those above or below, but little information transfer from those directly ahead. Information transfer in respect of changes in speed in the vertical plane is primarily focused on those directly ahead. Taken together, this suggests that krill adapt their heading based on those ahead and above or below, and adapt their speed based on those near neighbors who are directly ahead of them. The net effect of this is a pattern of alignment with those ahead and below and, to some extent, with those behind and above. One plausible explanation for the observed patterns is that the krill’s responses are governed to some extent by the hydrodynamics of krill swarms. Near neighbors produce a flow field as they swim, pushing water downwards and behind them, which emphasizes the importance of avoiding near neighbors who are above and ahead in particular (*15*). Decreasing speed and turning away when near neighbors are above may be an important part of this process. In addition, these strong responses in the vertical dimension may relate to krill predator-avoidance strategies or their mode of communication. For instance, many oceanic predators attack predominantly from above or below, rather from the side (*16,17*), while the positioning of bioluminescent photophores on the ventral surface mean that signaling between krill occurs predominantly in the vertical plane (*18*). The suggestion that the photophores act to counter-illuminate krill and make them less conspicuous to predators attacking from below potentially means that these points are interrelated (*18*).

Our analysis of krill interactions in three dimensions represents an important first step in understanding the principles underlying krill swarming behavior, however data from free-ranging krill in the Southern Ocean are urgently required to ground truth the laboratory-based observations presented here. Future work should examine the context-dependency of krill interaction rules in relation to oceanic currents, ambient light, temperature, predator avoidance and food availability. In addition, it would be valuable to examine the adaptive significance of the responsiveness of krill to near neighbors in the vertical dimension.

## Acknowledgments

We thank Norman Gaywood for his support for this project through his management of the Turing computational system at the University of New England, which was vital for the completion of this work.

## Author contributions

SK, JK, AJW conceived the idea; ALB, SK, MC, RK, AJW designed the project, ALB and AJW collected the data; TMS, JL, ALB, AJW analysed the data; ALB, AJW, TMS wrote the paper with contributions from all authors.

## Data and materials availability

All data will be deposited in Dryad upon acceptance.

## Supplementary Materials

Materials and Methods

Supplementary Text

Figures S1-S31

Tables S1-S2

References(*19-27*)

## References

1. Hamner WM, Hamner PP. Behavior of Antarctic krill (Euphausia superba): schooling, foraging, and antipredatory behavior. Canadian Journal of Fisheries and Aquatic Sciences. 2000;57:192–202.

2. Swadling K, Ritz D, Nicol S, Osborn J, Gurney L. Respiration rate and cost of swimming for Antarctic krill, Euphausia superba, in large groups in the laboratory. Marine Biology. 2005;146(6):1169–1175.

3. Kawaguchi S, King R, Meijers R, et al. An experimental aquarium for observing the schooling behavior of Antarctic krill (Euphausia superba). Deep-Sea Research Part Ii-Topical Studies in Oceanography. 2010;57(7-8):683–692.

4. Cox MJ, Warren JD, Demer DA, Cutter GR, Brierley AS. Three-dimensional observations of swarms of Antarctic krill (Euphausia superba) made using a multi-beam echosounder. Deep-Sea Research Part Ii-Topical Studies in Oceanography. 2010;57(7-8):508–518.

5. Hu F, Nie LJ, Fu SJ. Information Dynamics in the Interaction between a Prey and a Predator Fish. Entropy. 2015;17(10):7230–7241.

6. Tomaru T, Murakami H, Niizato T, et al. Information transfer in a swarm of soldier crabs. Artificial Life and Robotics. 2016;21(2):177–180.

7. Ward AJW, Schaerf TM, Burns ALJ, et al. Cohesion, order and information flow in the collective motion of mixed-species shoals. Royal Society Open Science. 2018;5(12):181132.

8. Herbert-Read JE, Perna A, Mann RP, Schaerf TM, Sumpter DJT, Ward AJW. Inferring the rules of interaction of shoaling fish. Proceedings of the National Academy of Sciences of the United States of America. 2011;108:18726–18731.

9. Katz Y, Tunstrom K, Ioannou CC, Huepe C, Couzin ID. Inferring the structure and dynamics of interactions in schooling fish. Proceedings of the National Academy of Sciences of the United States of America. 2011;108(46):18720–18725.

10. Ward AJW, Webster MM. Sociality: The Behavior of Group-Living Animals. Springer; 2016.

11. Ward AJW, Schaerf TM, Herbert-Read JE, Morrell LJ, Sumpter DJT, Webster MM. Local interactions and global properties of free-ranging stickleback shoals. Royal Society Open Science. 2017;4(7):170043.

12. Lukeman R, Li YX, Edelstein-Keshet L. Inferring individual rules from collective behavior. Proceedings of the National Academy of Sciences of the United States of America. 2010;107(28):12576–12580.

13. Attanasi A, Cavagna A, Del Castello L, et al. Collective Behavior without Collective Order in Wild Swarms of Midges. Plos Computational Biology. 2014;10(7).

14. Ballerini M, Cabibbo N, Candelier R, et al. Empirical investigation of starling flocks: a benchmark study in collective animal behavior. Animal Behavior. 2008;76:201–215.

15. Yen J, Brown J, Webster DR. Analysis of the flow field of the krill, Euphausia pacifica. Mar Freshw Behav Physiol. 2003;36(4):307–319.

16. Miller EJ, Potts JM, Cox MJ, et al. The characteristics of krill swarms in relation to aggregating Antarctic blue whales. Scientific Reports. 2019;9:13.

17. Silverman ED, Veit RR. Associations among Antarctic seabirds in mixed species feeding flocks. Ibis. 2001;143(1):51–62.

18. Grinnell AD, Narins PM, Awbrey FT, Hamner WM, Hamner PP. Eye photophore coordination and light-following in krill, euphausia-superba. Journal of Experimental Biology. 1988;134:61–77.

19. Kraskov, A., Stögbauer, H., & Grassberger, P. (2004). Estimating mutual information. Physical review E, 69(6), 066138.

20. Lizier, J. T. (2014). JIDT: An information-theoretic toolkit for studying the dynamics of complex systems. Frontiers in Robotics and AI, 1, 11.

21. Crosato, E., Jiang, L., Lecheval, V., Lizier, J. T., Wang, X. R., Tichit, P., … & Prokopenko, M. (2018). Informative and misinformative interactions in a school of fish. Swarm Intelligence, 12(4), 283–305.

22. Ballerini, M., Cabibbo, N., Candelier, R., Cavagna, A., Cisbani, E., Giardina, I., … Zdravkovic, V. (2008). Interaction ruling animal collective behavior depends on topological rather than metric distance: evidence from a field study. Proceedings of the National Academy of Sciences of the United States of America, 105(4), 1232–1237.

23. Calovi, D. S., Litchinko, A., Lecheval, V., Lopez, U., Pérez Escudero, A., Chaté, H., … Theraulaz, G. (2018). Disentangling and modeling interactions in fish with burst-and-coast swimming reveal distinct alignment and attraction behaviors. PLoS Computational Biology, 14(1), e1005933.

24. Couzin, I. D., Krause, J., James, R., Ruxton, G. D., & Franks, N. R. (2002). Collective memory and spatial sorting in animal groups. Journal of Theoretical Biology, 218(1), 1–11. doi:10.1006/yjtbi.3065

25. Heras, F. J. H., Romero-Ferrero, F., Hinz, R. C., & de Polavieja, G. G. (2019). Deep attention networks reveal the rules of collective motion in zebrafish. PLoS Computational Biology, 15(9), e1007354.

26. Schaerf, T. M., Dillingham, P. W., & Ward, A. J. W. (2017). The effects of external cues on individual and collective behavior of shoaling fish. Science Advances, 3, e1603201.

27. Fisher, N. I., Lewis, T., & Embleton, B. J. J. (1987). Statistical analysis of spherical data: Cambridge University Press.

