## Supplementary Information for "Self-organization and information transfer in Antarctic krill swarms"

**This PDF file includes:**

Materials and Methods

Supplementary Text

Figs. S1 to S31

Tables S1 to S2

Materials and Methods

Study species

Antarctic krill were collected by midwater trawl from the Southern Ocean during the 2016/17 field season. The krill used in this study (average length ~40mm) were kept at the Australian Antarctic Division’s marine research aquarium at Kingston, Tasmania, in an 1860L cylindrical tank (see *3*).

Filming and camera calibration

Two Gopro™ Hero 6 cameras were used for filming at a rate of 30 frames per second for at least 30 minutes at a time. Cameras were fixed on an aluminium frame and submerged approximately 50cm beneath the surface of the water. The tanks were covered with white corflute for the duration of filming to minimize any disturbance by light or people walking by. In order to facilitate tracking, Gopros™ were positioned to film against this white background – i.e. pointing vertically upwards.

In order to calibrate the cameras, a black and white printed grid was moved through the field of view of both cameras while submerged in each tank. Videos were then calibrated using the Stereo Camera Calibrator application in MATLAB (https://au.mathworks.com/help/vision/ug/stereo-camera-calibrator-app.html). Using this tool we were able to determine the intrinsic and extrinsic parameters of each camera and the distortion coefficients which would allow us to convert our images to 3-dimensions. Calibration accuracy and reprojection error were set manually to less than 1 pixel, so that the images on each camera matched to within 1 pixel. The mean reprojection error was 0.53 pixels. See Figure S1 for images relating to the calibration process.

Tracking

Individual krill were tracked manually in ImageJ from 10-second clips, with these clips further subdivided into smaller duration sections. Clips were cut to 10 seconds or less as this was typically the length of time a single krill would spend in the field of view of both cameras. Coordinate data were then imported into MATLAB where matched pairs of (*x, y*) coordinates were first corrected to take into account effects of camera distortion using the *undistortpoints* function, and then converted to three dimensions with (*x, y, z*) coordinates given in millimetres using the *triangulate* function. In the (*x*, *y*, *z*) coordinate system, the *z*-coordinate corresponded to the direction perpendicular to the Earth’s surface, with the *x* and *y* coordinates describing displacements in the horizontal. For this study we focussed our analysis on data derived from 10 clips of durations from 50 to 788 frames, with tracks obtained for 20 to 55 krill for each of these clips.

**Supplementary Text:**

Transfer entropy

We used the Kraskov, Stögbauer and Grassberger estimator (*19*) from the Java Information Dynamics Toolkit (JIDT) open-source software (*20*) via the demos/octave/Flocking scripts, to quantify information flow. Using time-series data (*x,y,z*), we measured transfer entropy (in nats) based on both changes in heading direction and speed, whereby greater levels of TE suggest greater potential information transfer from one individual to another. Specifically, transfer entropy measures the information held about the target variable (in this case the change in target krill heading direction or speed) by the source variable, or the heading direction or speed of a source krill relative to the heading direction or speed of a target krill. As per (*21*), all source-target pairs within range across every frame of all 10 clips were used to create samples for the whole dataset, meaning that the TE measured for a clip is an ensemble average of the representative pairwise source-target interaction for all krill pairs at every frame in the trial.

TE optimization

Before calculating the Transfer Entropy results, we ran two optimizations for the source-target lag, embedding dimension k and delay tau (similar to that detailed in *21*). Firstly, with a pair range set at ‘infinite’ so that all individuals were considered, and then after determining that individuals almost exclusively interact within 200mm we set this as our maximum pair range (see optimized parameters Table S1):

Estimating rules of interaction in three spatial dimensions

Here, rules of interaction refer to the set of rules for how an individual adjusts its velocity as a function of: the relative coordinates of its groupmates, the individual’s current behavior, for example the current velocity of the individual, and the behavior of group mates, which might also include their velocities. Although the initial studies that inferred the presence of interaction rules based on local group structure were conducted using data with three spatial dimensions (in studies of starling murmurations, such as that in (*22*)), many subsequent studies have focused on inferring the presence or form of local interactions for movements in two dimensions *(8, 9, 12, 23-26*) . To the best of our knowledge, there have been no studies to date that have explicitly determined how individuals adjust their velocity as a function of the relative coordinates of group mates in three dimensions. Here we build on previous work, particularly that of *(8, 9, 12, 22, 26*), to develop a method for identifying how an individual adjusts its velocity as a function of the relative coordinates of its group mates when moving in three spatial dimensions.

We write the position of individual *i* at some discrete time *t* in three dimensions as
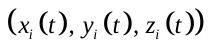
, where *x* and *y* represent the individual’s position in the horizontal direction, and *z* indicates the individual’s position in the vertical. Discrete time steps are separated by equal durations of
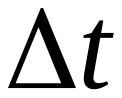
. Throughout our calculations, variables in bold type represent vectors, we write the unit vectors parallel to the positive *x*-, *y*-, and *z*-axes as **i**, **j**, and **k** respectively, an expression of the form
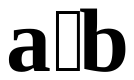
 denotes the scalar dot product of the vectors **a** and **b**, and an expression of the form
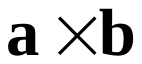
denotes the vector cross product of the vectors **a** and **b**. The fundamental calculations described below are aimed at describing changes in the components of an individual’s velocity via changes in speed (the magnitude of velocity) and changes in direction of motion via two angles (
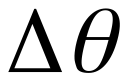
, which describes a rotation about the *z*-axis/in the *xy*-plane, and
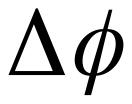
, which describes a rotation up or down relative to the *xy*-plane).

Estimates for velocity, speed, changes in speed, and turning speed

Analogous to previous work in two dimensions (*26*), we first smooth each individual’s trajectory by applying the Savitzky-Golay method with degree 2 and span 5 (default settings as applied by MATLAB’s *smooth* function) to the time series of *x*-, *y*-, and *z*-coordinates. We then estimate the components of an individual’s velocity, in the *x*- , *y*-, and *z*-directions respectively, at time *t* via the forward difference approximations

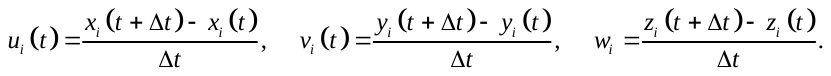

Individual *i*'s approximate velocity vector at time *t* is thus
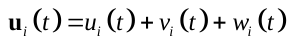
. The corresponding approximate speed of individual *i* at time *t* is:

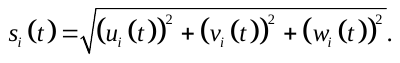

It follows immediately that the unit vector pointing in the direction of motion of individual *i* is:

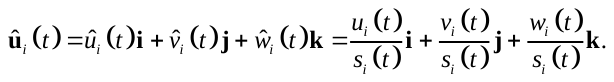

We then estimate the change in speed of individual *i* at time *t* via:

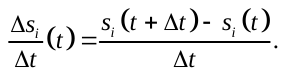

The quantity above captures both the direction and magnitude of speed changes, with positive values of
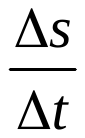
 indicating increases in speed, and negative values of
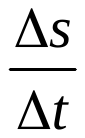
 indicating decreases in speed.

To examine if individuals tended to match their speed differently with group mates in different relative positions, we calculated the absolute differences in speed between all pairs of krill, *i* and *j*, for all discrete times *t*:
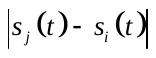
.

We estimated individual turning speeds at given times via:

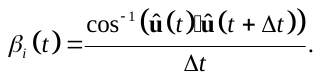

Estimates for changes in direction

We determine changes in direction by comparing the relative directions of
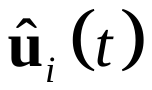
 and
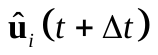
. To aid in these calculations, we adopted the same convention for a consistent frame of reference for individuals as that used by (*22*); in the calculations that follow we adjust coordinate frames such that individual *i* is located at the origin of the coordinate system
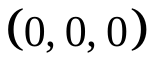
, individual *i*’s velocity vector at time *t* is parallel to the positive *x*-axis, and the component of gravity perpendicular to the individual’s velocity vector at time *t* is parallel to the negative *z*-axis. The unit vector in the direction of gravity is
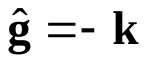
. The component of the unit vector in the direction of gravity parallel to
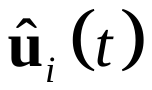
 can thus be written as
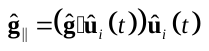
. It therefore follows that the component of
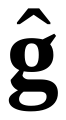
perpendicular to
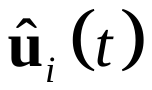
 is
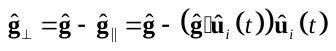
. The individual’s occupancy of the origin is implicit in our calculations for changes in direction, and we use a sequence of two rotations about the location of the individual to align
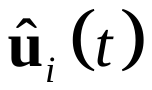
 and
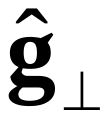
 with the positive *x*- and negative *z*-axes respectively.

First, we determine the angle,
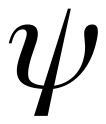
, between the projection of
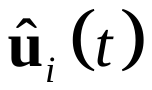
 onto the *xy*-plane (
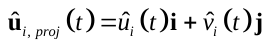
) and the positive *x*-axis. A computationally convenient way to do this is via
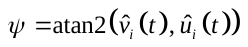
, where atan2 is a common computational application of the inverse tangent function that correctly identifies both the sign and magnitude of
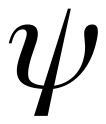
 such that
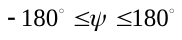
. We then construct the following rotation matrix to rotate coordinates through an angle of
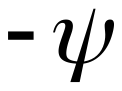
 about the *z*-axis (such that
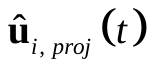
is perfectly aligned with the positive *x*-axis):

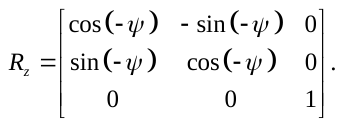

We use the rotation matrix,
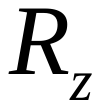
, to rotate the components of
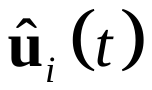
 and
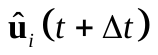
 about the *z*-axis to obtain:

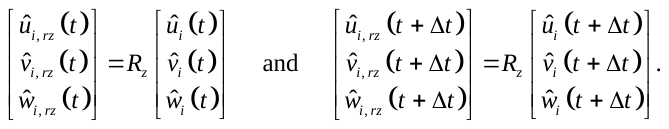

Next, we determine the angle made between
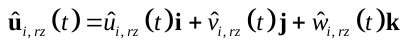
 and the *x*-axis (in the *xz*-plane); this angle is the same as the angle that
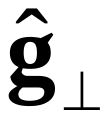
 makes with the negative *z*-axis after rotation via
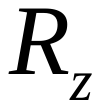
, and can be determined via
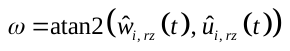
. With this angle, we construct a second rotation matrix,
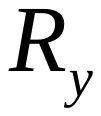
, that rotates points through an angle of
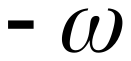
 about the *y*-axis, such that what was originally the vector
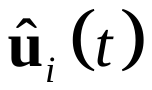
 will be completely aligned with the positive *x*-axis, and as a consequence
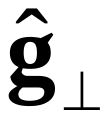
 will align with the negative *z*-axis. The appropriate rotation matrix is:

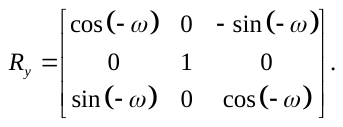

We then rotate
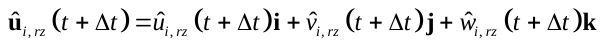
using
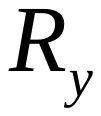
 via:

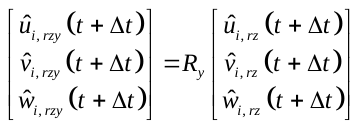
.

The above calculation ultimately gives the components of
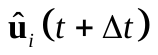
 relative to
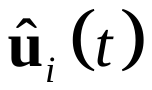
 in a three dimensional rectangular coordinate system; that is the change in direction of individual *i* from observation times *t* to
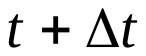
. The change in direction can be described via a combination of two angles,
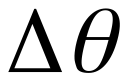
 and
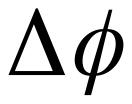
, as mentioned above, but for computational convenience we retain the rectangular coordinate representation of the change in angle (described by
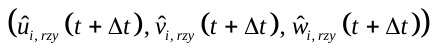
) until later in our calculations. Using analogous calculations, we also determine the components of
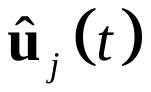
 in the reference frame where
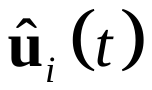
 is aligned with the positive *x*-axis, so as to examine the directions of motion of group members relative to an individual at the origin of our relative coordinate system; we denote the components of the direction of motion of individual *j* relative to individual *i* at time *t* using
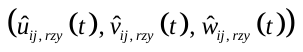
.

Relative coordinates, aggregation and binning of data, and averaging of binned values

The next stage in calculations is to determine the coordinates of group mates relative to each individual *i* (for each discrete time *t*) in a consistent coordinate frame where individual *i* is located at the origin, individual *i*’s velocity is parallel to the positive *x*-axis, and the component of gravity perpendicular to individual *i*'s velocity is aligned with the negative *z*-axis. Unlike the calculations above for determining changes in direction, there must be an explicit change of coordinates to place individual *i* (the focal individual) at the origin. An appropriate change of coordinates is achieved by subtracting the coordinates of individual *i* from that of their group mate to give:

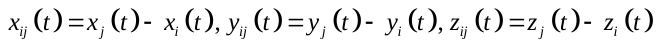
.

The same process that was used to rotate components of velocity to determine relative changes in direction is then applied to rotate the coordinates of individual *j* relative to individual *i*. We write the coordinates of individual *j* relative to individual *i*, such that
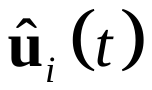
 points in the same direction as the unit vector
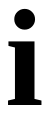
 and
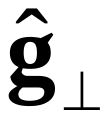
 points in the same direction as
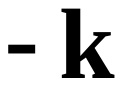
, as
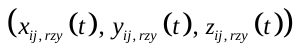
.

We divide a cubic domain centred on the focal individual into a set of
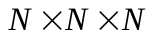
 overlapping cubic bins. The length scale of the cubic domain depends on the focus of a given study (for example short range, or longer range, interactions), but based on our previous experience with similar calculations in two dimensions, we have experimented with the domain extending from about 4 to 10 body lengths in the *x*, *y*, and *z* directions. (Based on our results, our choice of length-scale seems to have been enough to capture the presence of repulsion- and attraction-like changes in speed and direction.) The bins are used to store a behavioral measure of interest, such as the speed of individuals, changes in speed or direction, and absolute speed differences between individuals and their partners.

By means of example, we describe the binning process for individual speeds here, but the process is the same for all other measures. We deposit the speed of the focal individual,
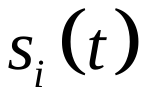
 , into all bins containing the relative coordinates of their group mates,
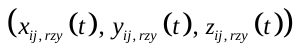
, aggregating data from all partners
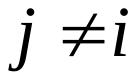
, for all discrete times, *t*, treating all individuals *i* in turn as the focal individual, and pooling data across multiple sets of observations. Once this process is complete, we then determine the mean speed value in each bin, with the result being an approximation to the mean speed of an individual as a function of the relative
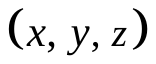
 coordinates of their partners, in a frame of reference where the direction of motion of the individual is aligned with the positive *x*-axis. Similar calculations are used to examine mean changes in speed of individuals, speed differences, and heading differences, as a function of relative partner coordinates, with simple means taken of the values collated in each bin. In addition, it is possible to construct a local density plot that describes the relative frequency that partners occupy given relative
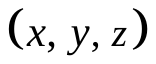
 coordinates by first counting the number of elements in each bin, and then dividing these bin-by-bin counts by the total count across all bins.

More complex is the examination of changes in direction. For these calculations, we deposit the components of the unit vectors
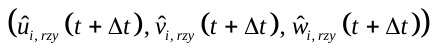
 to bins, as above. We then apply a standard method from circular and spherical statistics to determine the mean direction described by the vectors in each bin (*27*). We form the resultant vector by addition of all unit vectors in a given bin. Writing
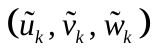
 as the components of the *k*th of *m* unit vectors stored in a particular bin, the resultant vector will then have components:

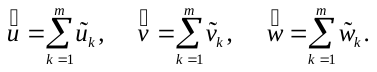

This resultant vector in fact points in the mean of the directions contained in a given bin, and we determine the corresponding unit vector pointing in the same direction by dividing the above components by the magnitude of the vector, given by
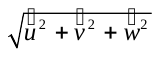
. Writing the components of a resulting unit vector for a given bin as
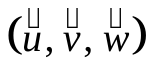
, we then rewrite the direction indicated by the vector in terms of the angle made between the projection of the vector onto the *xy*-plane and the positive *x*-axis, denoted
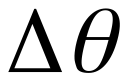
, and the angle made between the vector and the *xy*-plane, denoted
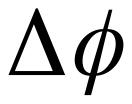
. In practice, we determine these angles via:

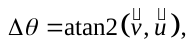

and,

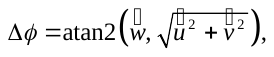

with the resulting change in direction over time described by
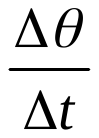
 and
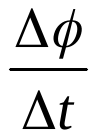
. Our choice for measuring the angle
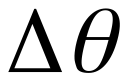
 is consistent with the standard mathematical convention for describing an angle in the *xy-*plane in spherical coordinate systems, and is equivalent to describing longitude on the surface of a sphere. When viewed from above, positive values of
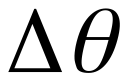
 coincide with anticlockwise rotations about the *z*-axis, and negative values of
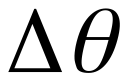
 coincide with clockwise rotations, with
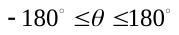
. This is also analogous to our measure for changes in direction in the plane when examining interactions in two-dimensions (*8, 26*). As defined above,
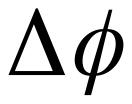
 describes latitude on the surface of a sphere, with positive values of
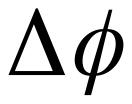
 coinciding with upward turns, and negative values of
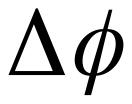
 coinciding with downward turns.
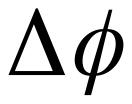
 is constrained such that
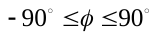
. Our convention for constructing
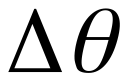
 and
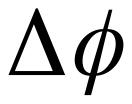
 is illustrated in Figure S2.

The calculations that we performed to examine relative directions of motion of group mates as a function of their relative coordinates are initially very similar to those for examining the turning components of velocity. We deposit the components
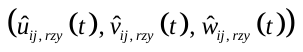
 determined in earlier calculations into all the bins containing
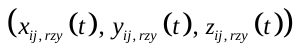
. We then sum the vectors contained in each bin, and determine the unit vector pointing in the same direction as the resultant, writing the unit vector as
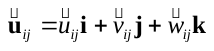
. (This unit vector points in the mean direction of the angles stored in a given bin.) The angle between the direction of the unit vector
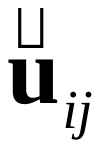
 and that of the focal individual (aligned with the positive *x*-axis) can then be determined via
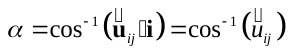
.

The above calculations can be modified to reduce the number of independent variables that are taken into account. For example, it is possible to examine the mean speed of individuals as function of only the relative
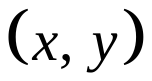
, , or coordinates of group mates, with the associated functions referred to as projections onto the *xy*-, *xz*-, or *yz*-planes respectively. Such projections can be constructed by either performing the binning process in two dimensions only, or by aggregating data across a particular dimension after binning has been performed in three dimensions (which is the approach that we’ve applied for this study).

Using additional RAM to increase calculation speed

Two of the most computationally time-consuming elements of the process of fitting the types of functions described in the previous section are the pairwise determination of the relative coordinates of group mates, and the triple nested loop over the array of cubic bins. We reorganised components of our calculations to try to avoid repeating these elements.

For a group of *n* individuals, a total of pairwise comparisons are required to determine the coordinates of each group mate relative to that of a focal individual. There will not usually be any symmetry in the coordinates of individual *j* relative to individual *i* and the coordinates of individual *i* relative to individual *j* due to the fact that the relative coordinate system takes into account the direction of motion of the focal individual (not just their absolute position in space). Given the operation count for determining relative coordinates, it is desirable to try to avoid repeating these calculations. The approach that we have applied is to perform a once and for all set of calculations to determine the relative coordinates of group mates compared to focal individuals for all discrete time steps in a given observation set. This calculation is performed before entering the loops used to fill the array of cubic bins. We store the coordinates of group mates relative to their partners using three matrices (one matrix for *x*-coordinates, one matrix for *y*-coordinates, and one for *z*-coordinates), where is the number of output times; since the relative coordinate system uses the velocity of the focal individual as a reference, relative coordinates can only be determined consistently for time steps. The first *n* rows of these matrices are filled with the coordinates of group mates relative to individual 1, the next *n* rows with the coordinates of group mates relative to individual 2, and so on, up to the *n*th individual. Any row corresponding to the coordinates of an individual *i* relative to itself is filled with not-a-number placeholders. Alongside the matrices of relative coordinates, we construct analogous matrices that contain the components of group mates’ directions of motion relative to that of focal individuals, and a matrix that records the absolute speed difference between focal individuals and their partners for all time steps. We also construct additional matrices that contain multiple copies of the time series of the speed, changes in speed, and components of direction changes of focal individuals, with the same dimensions as the matrices containing relative coordinates. The matrix containing instantaneous speed estimates is constructed so that the first *n* rows contains copies of the time series of observed speeds for individual 1, the second *n* rows contains copies of the time series of observed speeds for individual 2, all the way up to individual *n*, with the same process then applied for storing changes in speed, and the components of direction changes. For subsequent calculations, only data up to time step is then used, corresponding to the number of time steps for which there are estimates for changes in velocity.

After storing relative coordinates, components of velocity, speed differences, individual speeds, changes in speed, and components of direction changes in the above matrices/lookup tables, the triple nested loop over the array of bins is then entered. For each bin, the indices of all elements that correspond to points contained within that bin are identified from the matrices containing relative *x*, *y*, and *z* coordinates. These indices are then used to extract and bin the corresponding components of relative directions of motion, speed differences, speed of focal individuals, changes in speed, and components of direction changes, which are then stored in the bins (in practice we store binned data using cell arrays in MATLAB, and we exploit MATLAB’s linear indexing of arrays to simplify our code). In addition, the number of elements to be included in each bin is counted in order to construct an approximation to the relative frequency that individuals occupy given relative coordinates, as noted above.

The approach above guarantees that only one search operation must be applied to identify all elements that should be placed in a given bin, across all pairwise relations between individuals, and for all time steps. In practice we’ve found that the above ordering of calculations results in an approximate threefold increase in calculation speed, as compared to earlier versions of our codes where binning was repeated for each pair of individuals. The increased speed does however come at the cost of an increase in memory usage. For example, a group of 50 individuals observed over a duration of 1000 time steps will require twelve 2500 × 998 matrices to store the measures discussed throughout this supplementary material in lookup form. If each matrix stores standard double precision elements, then the overall memory requirement for this example case is 239520000 bytes (239.52 megabytes), which is well within the RAM capacity of most modern computers. However, care must still be taken, as the storage requirement will increase proportional to the number of individuals squared, and these lookup matrices are not the only arrays that are stored during calculations (for example, the potentially large cell arrays containing binned data also require a portion of the available computational memory).

Results

Figure S3 contains histograms of the observed speeds, changes in speed, and turning speeds of the krill (aggregated from all 10 sets of trajectories).

We performed calculations over two different scales and resolutions. To examine the relative frequency that neighbors were encountered at given relative (*x, y, z*) coordinates and relative alignment with near neighbors in greater detail close to the focal individual, we performed the calculations described in earlier sections over the volume where -200 ≤ *x, y, z* < 200 mm, divided into overlapping cubic bins of side length 20 mm (approximately half a body length for the krill), with the closest sides of adjacent bins separated by 5 mm. We also performed calculations over the larger volume where -320 ≤ *x, y, z* < 320 mm, divided into cubic bins of side length 40 mm, with the closest sides of adjacent bins separated by 10 mm.

Figures S4 to S27 include additional graphs illustrating the statistical density of neighbors, relative alignment, mean individual speed, mean change in speed, and the components of mean direction changes as a function of the relative coordinates of neighboring krill. Our data analysis suggested that the turning response of the krill was dependent on the octant occupied by their neighbors, when examined in the relative coordinate system where the turning krill was located at the origin, with its current direction of motion aligned with the positive ­*x*-axis, and the component of gravity perpendicular to the turning krill was aligned with the negative *z*-axis. We summarise the broad observed octant by octant turning behavior in Table S2.

Figures S28 to S31 illustrate the mean absolute difference in speed between focal individuals and their neighbors, as a function of the relative coordinates of the neighboring krill. Based on an examination of these plots, there does not seem to be a clear pattern of matching speeds with neighbors over the plotted domain.

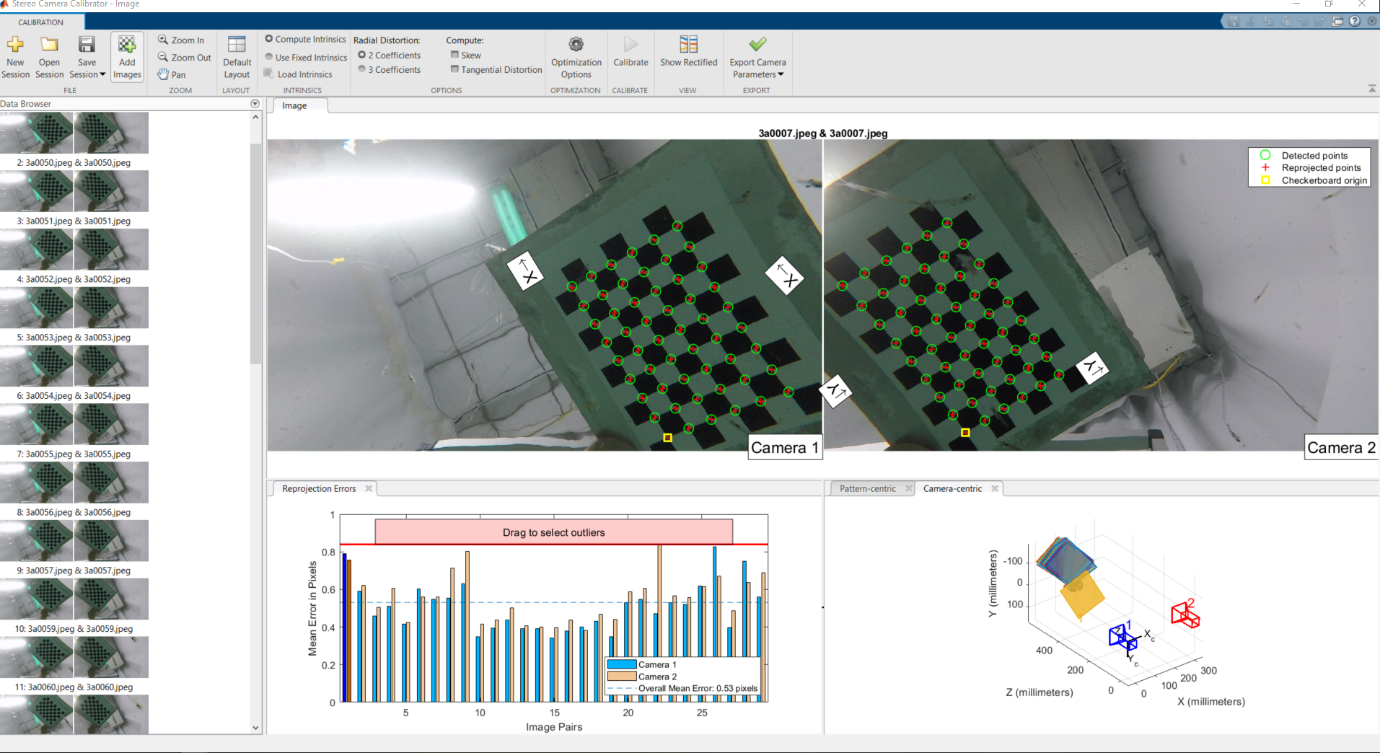

**FIGURE S1:** Snapshot of the calibration process applied in MATLAB.

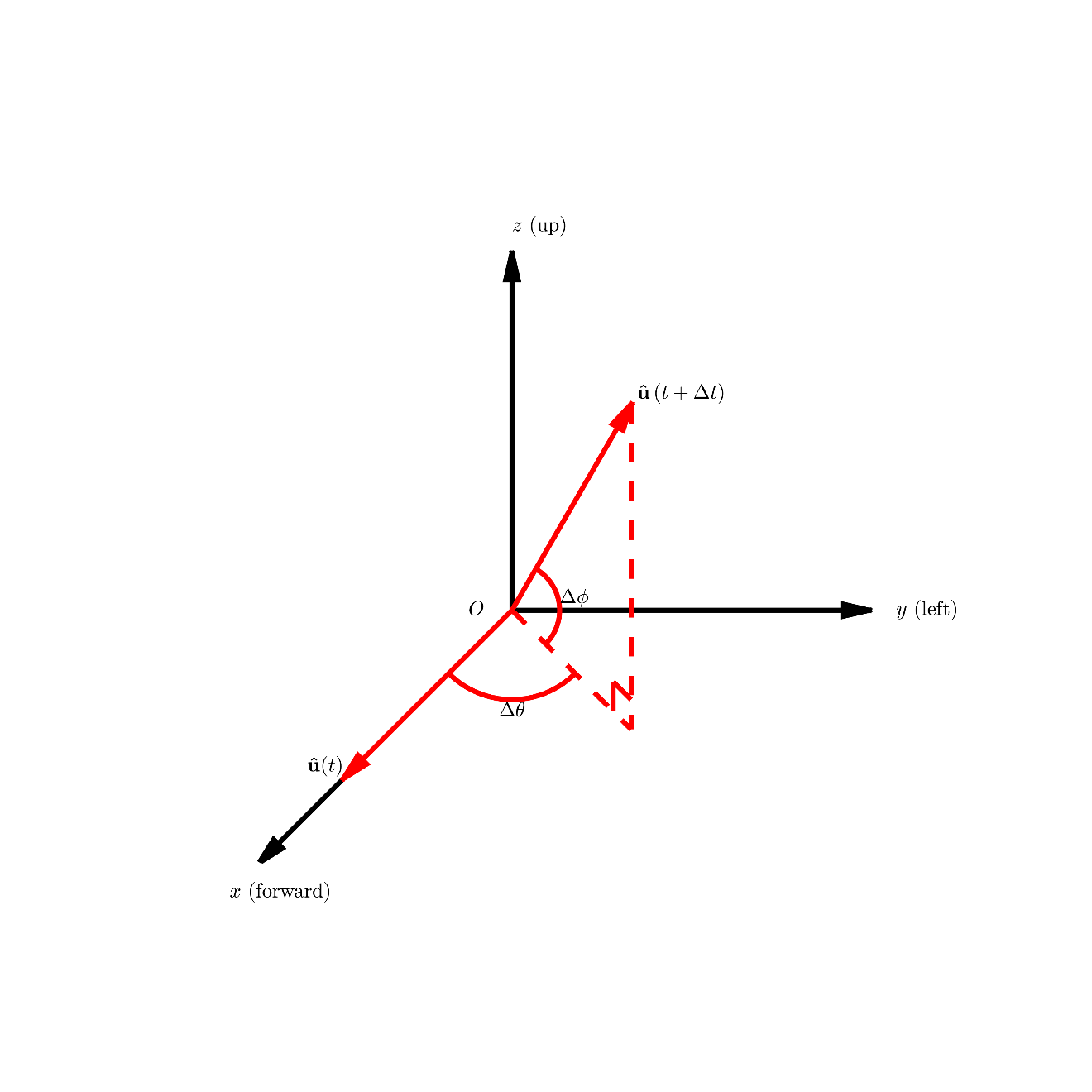

**FIGURE S2:** Our convention for identifying the components of velocity that describe the change in direction of an individual between consecutive video frames at times and . The direction of motion of a focal individual at time , denoted , is moved to a consistent frame of reference such that is aligned with the positive -axis, and the component of gravity perpendicular to is aligned with the negative *z*-axis. The direction of motion of the individual at time , , is then transformed to the same coordinate system, and the differences in direction are then measured in terms of (in the *xy*-plane) and (measured upwards or downwards relative to the *xy*-plane), as illustrated.

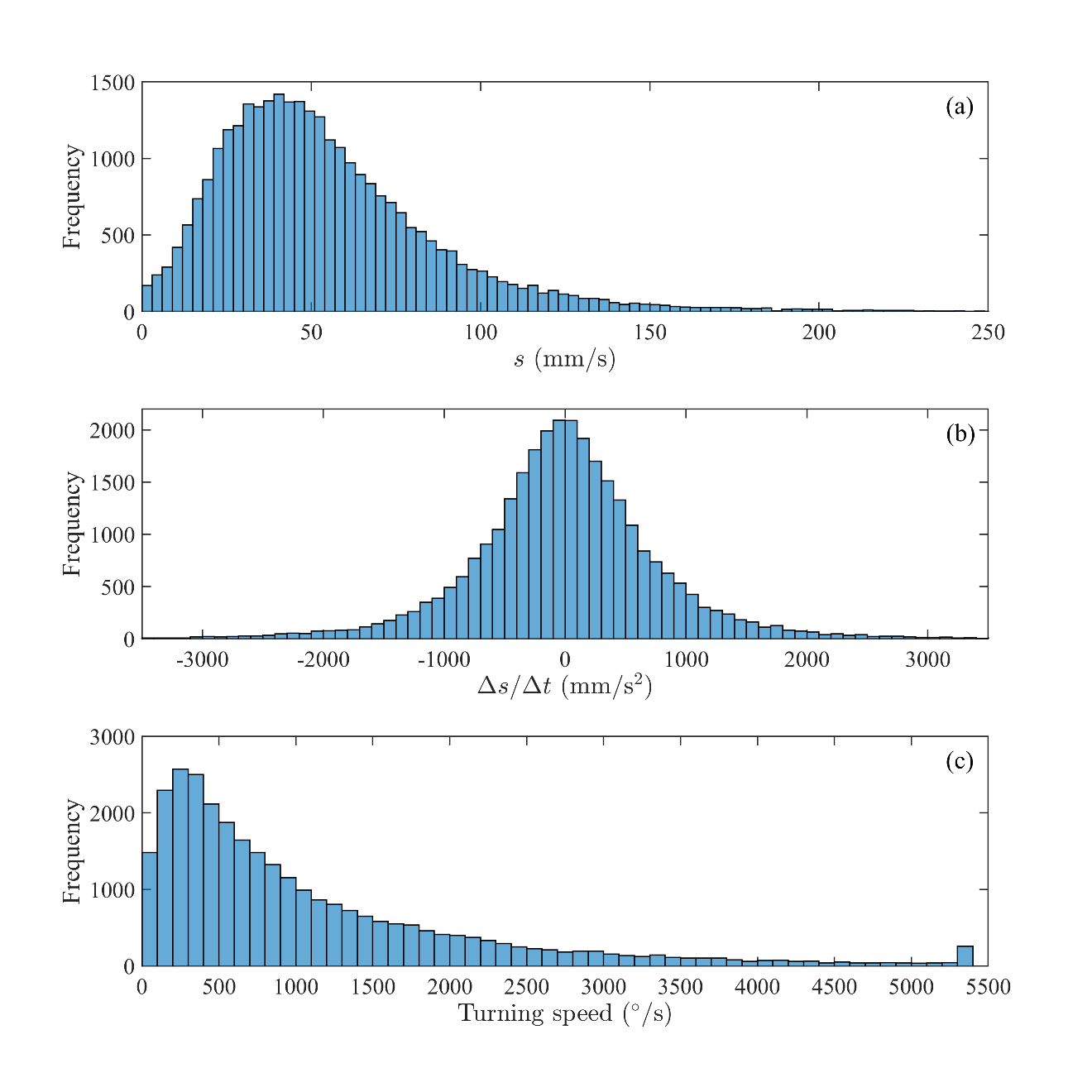

**FIGURE S3:** Histograms of the observed speeds, (a), changes in speed over time, (b), and turning speeds, (c), of individual krill for all tracked data. Note that at a frame rate of 30 frames per second, the maximum detectable turning speed is 5400 degrees per second (corresponding to a 180° change in direction between two video frames).

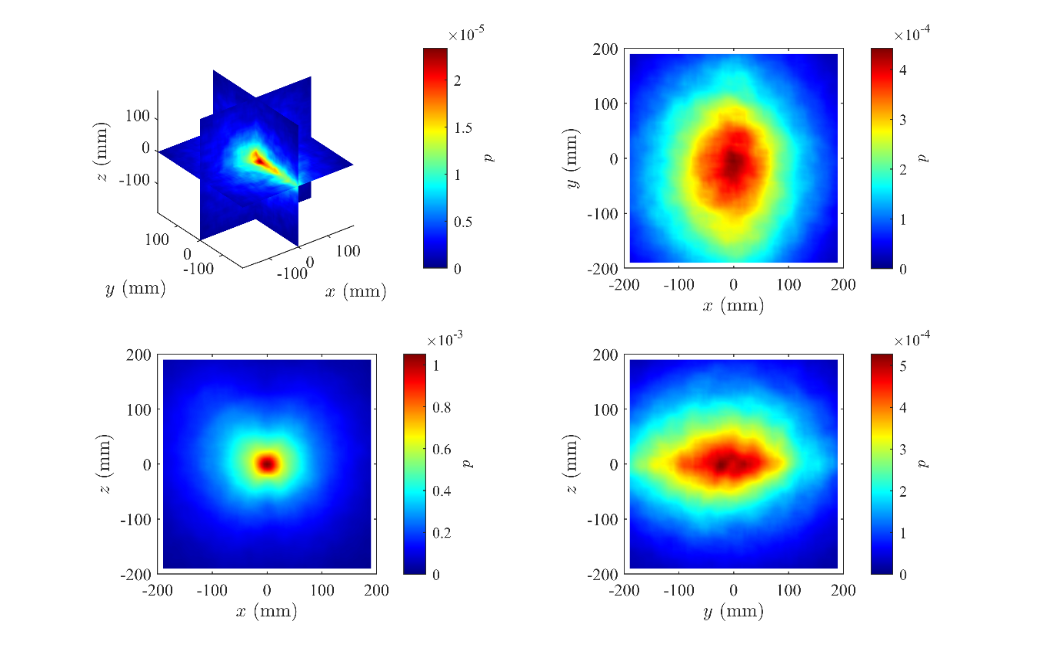

**FIGURE S4:** The statistical density of neighboring krill as a function of their (*x*, *y*, *z*) coordinates relative to a focal krill at (0, 0, 0), rendered in the planes *x* = 0, *y* = 0, and *z* = 0 (top left panel). The direction of motion of the focal individual is aligned with the positive *x*-axis, and the component of gravity perpendicular to the individual’s direction of motion is aligned with the negative *z*-axis. The other panels illustrate the density of neighboring krill as a function of relative (*x*, *y*) (top right panel), relative (*x*, *z*) (bottom left panel), and relative (*y, z*) (bottom right panel) coordinates only.

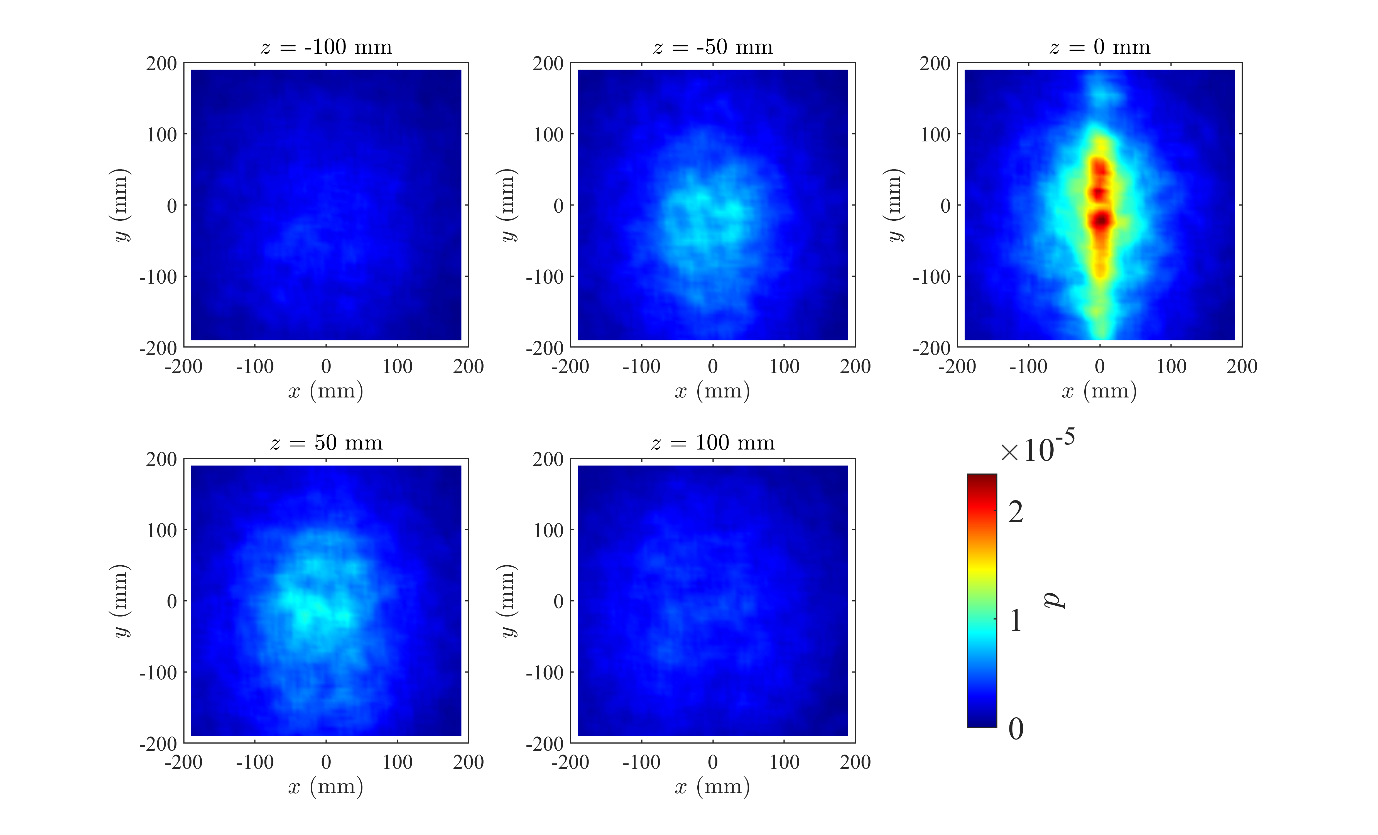

**FIGURE S5:** The statistical density of neighboring krill as a function of their (*x*, *y*, *z*) coordinates relative to a focal krill at (0, 0, 0), in the planes where *z* = -100 mm, *z* = -50 mm, *z* = 0 mm, *z* = 50 mm, and *z* = 100 mm. Positive *z* corresponds to regions above the focal individual, whereas negative *z* corresponds to regions below the focal individual.

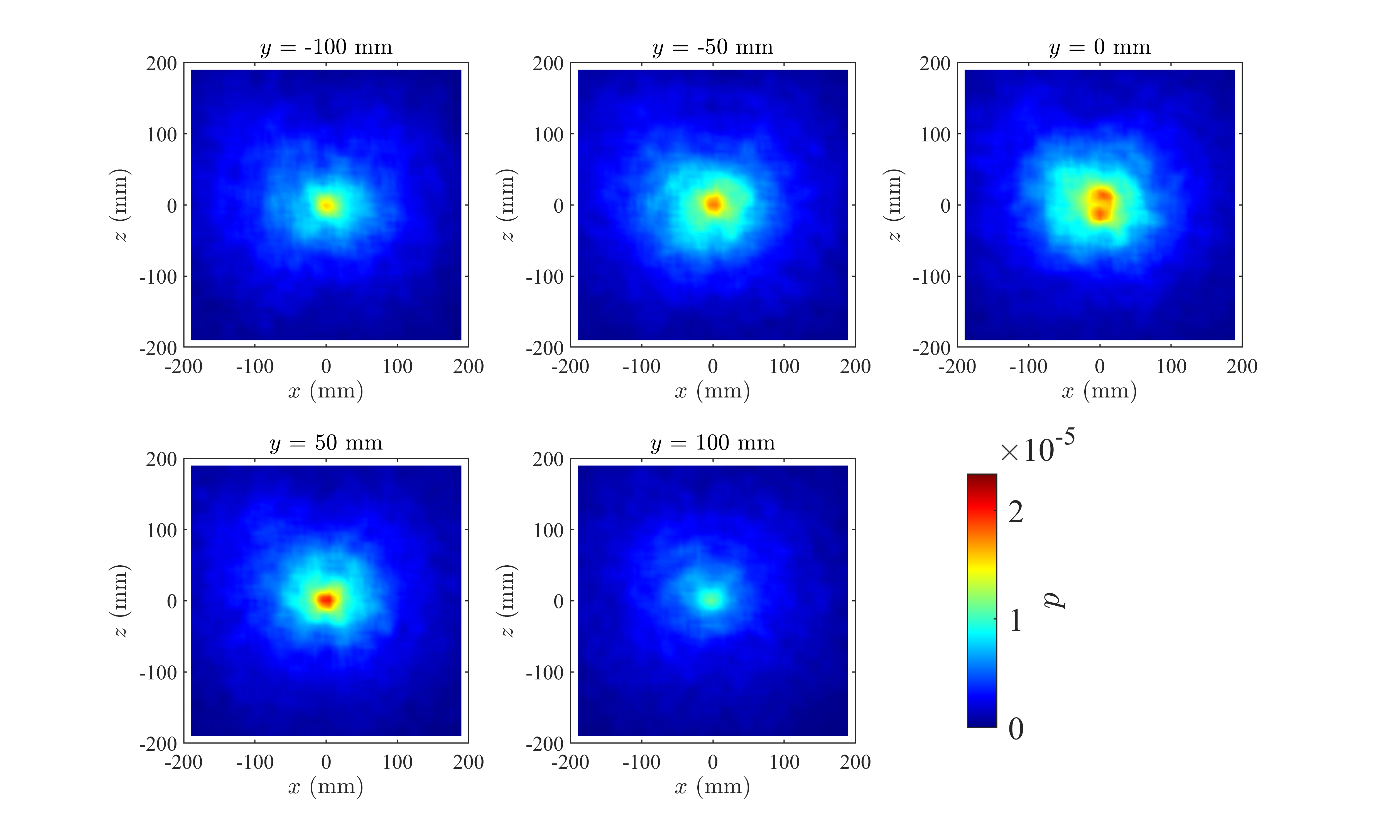

**FIGURE S6:** The statistical density of neighboring krill as a function of their (*x*, *y*, *z*) coordinates relative to a focal krill at (0, 0, 0), in the planes where *y* = -100 mm, *y* = -50 mm, *y* = 0 mm, *y* = 50 mm, and *y* = 100 mm. Positive *y* corresponds to regions to the left of the focal individual, whereas negative *y* corresponds to regions to the right of the focal individual.

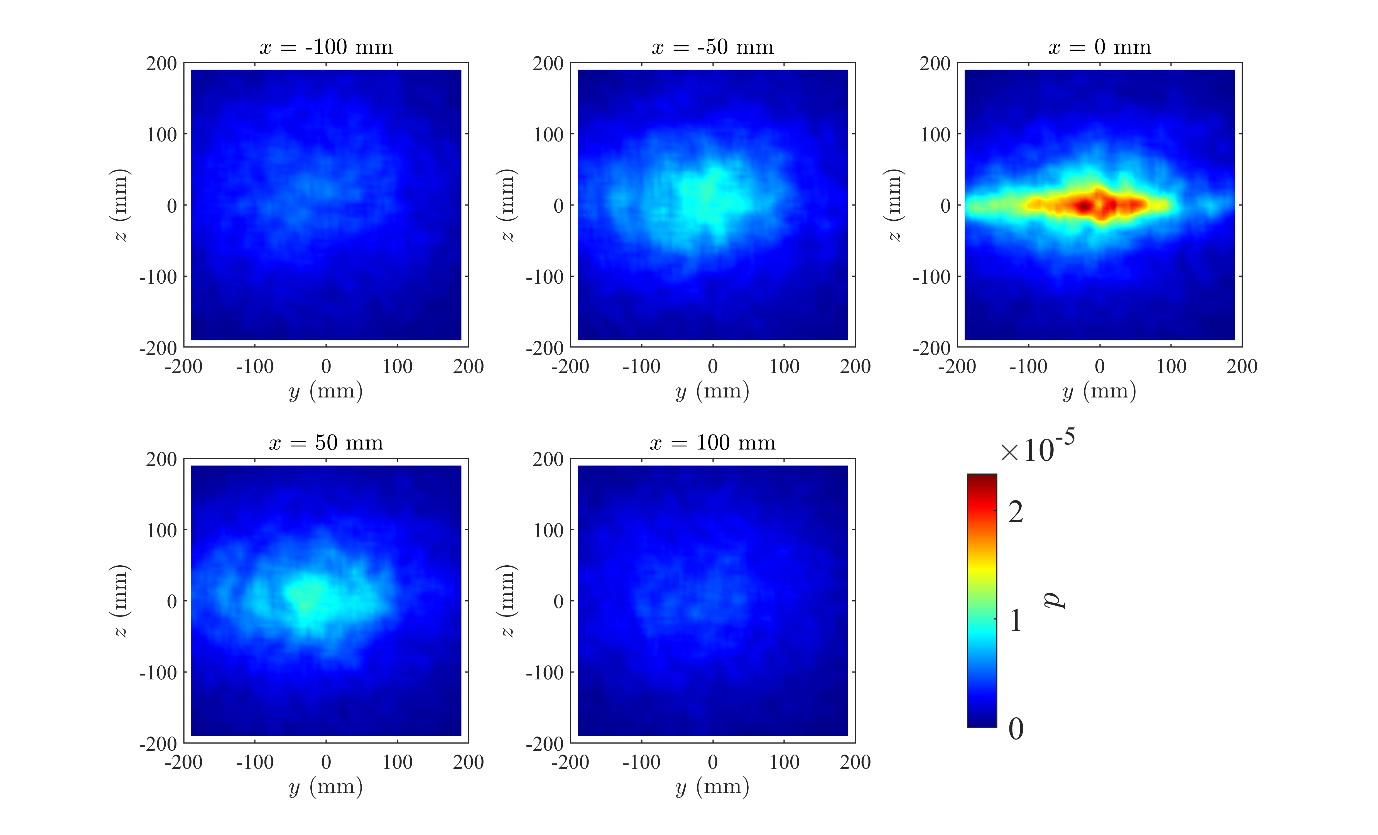

**FIGURE S7:** The statistical density of neighboring krill as a function of their (*x*, *y*, *z*) coordinates relative to a focal krill at (0, 0, 0), in the planes where *x* = -100 mm, *x* = -50 mm, *x* = 0 mm, *x* = 50 mm, and *x* = 100 mm. Positive *x* corresponds to regions in front of the focal individual, whereas negative *x* corresponds to regions behind the focal individual.

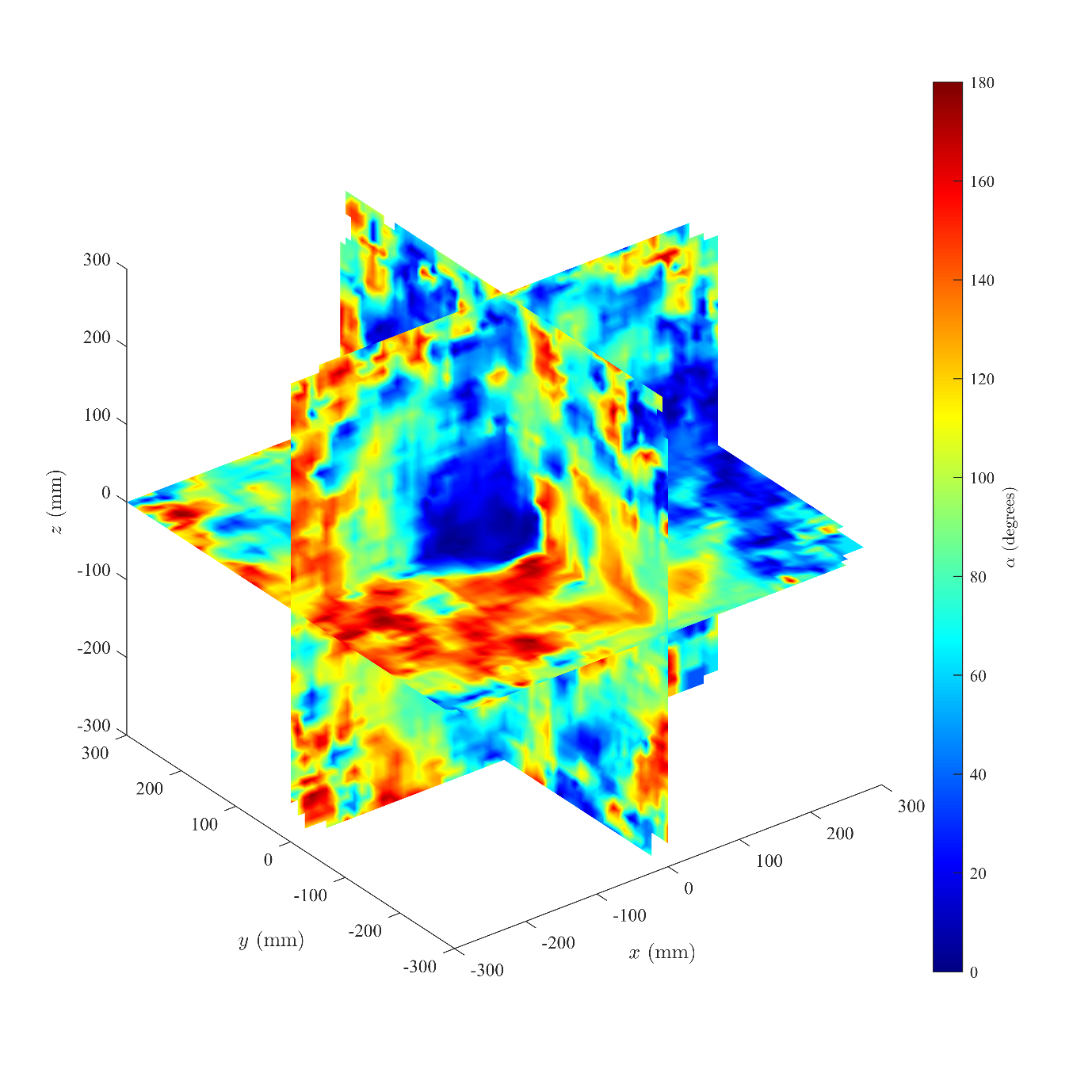

**FIGURE S8:** Mean angular difference in travelling direction, α, measured in of neighboring krill relative to a focal individual positioned at (0, 0, 0), with the focal individual travelling parallel to the positive *x*-axis, and the component of gravity perpendicular to the focal individual aligned with the negative *z*-axis (rendered in the planes *x* = 0, *y* = 0, and *z* = 0).

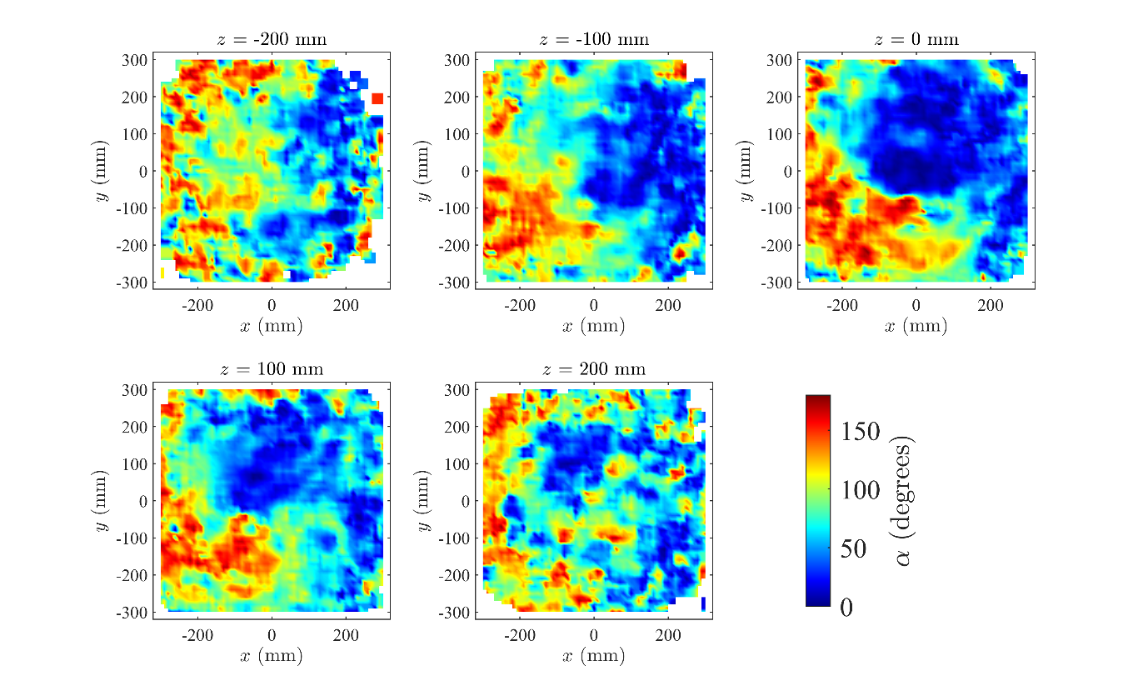

**FIGURE S9:** The mean angular difference in travelling direction of neighboring krill as a function of their (*x*, *y*, *z*) coordinates relative to a focal krill at (0, 0, 0), in the planes where *z* = -200 mm, *z* = -100 mm, *z* = 0 mm, *z* = 100 mm, and *z* = 200 mm. Positive *z* corresponds to regions above the focal individual, whereas negative *z* corresponds to regions below the focal individual.

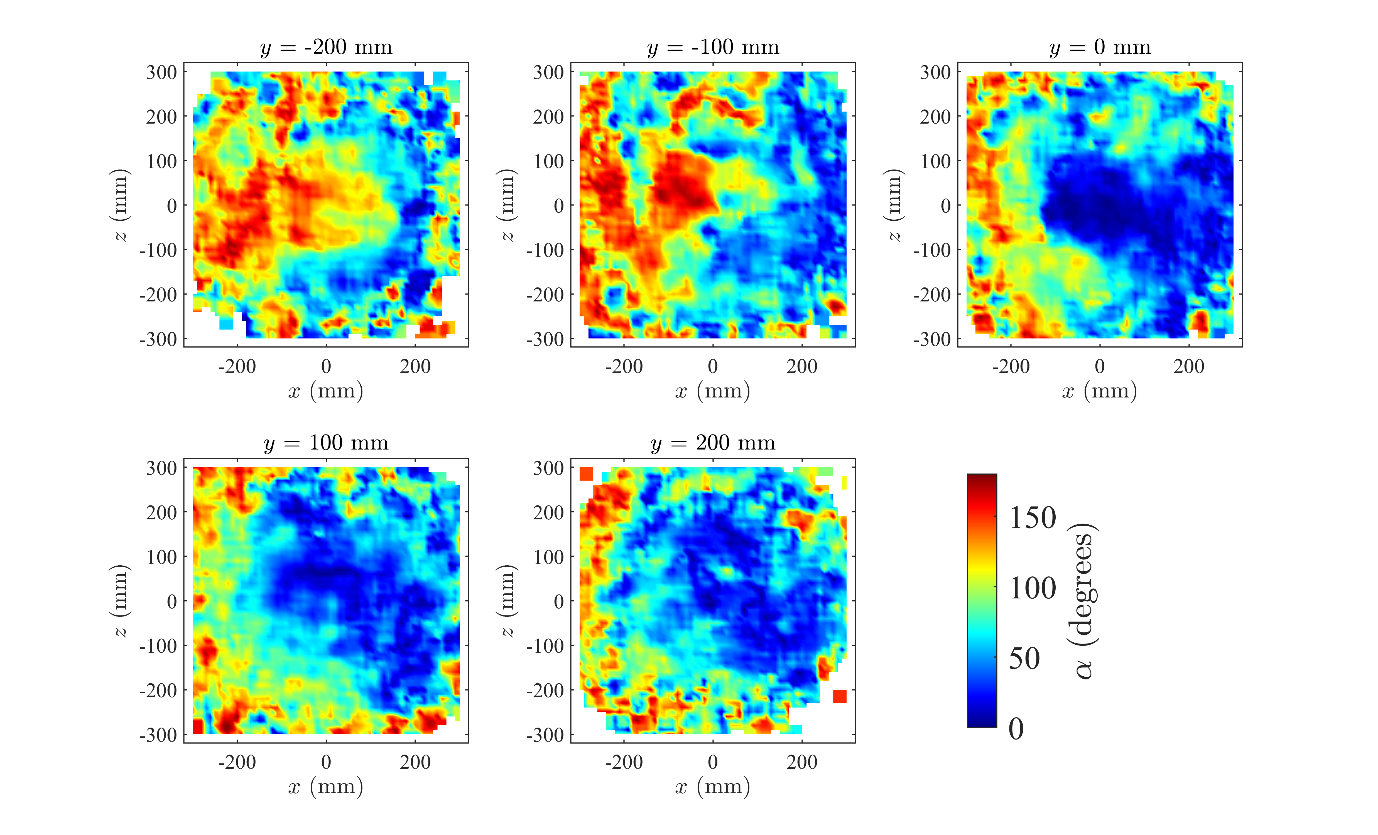

**FIGURE S10:** The mean angular difference in travelling direction of neighboring krill as a function of their (*x*, *y*, *z*) coordinates relative to a focal krill at (0, 0, 0), in the planes where *y* = -200 mm, *y* = -100 mm, *y* = 0 mm, *y* = 100 mm, and *y* = 200 mm. Positive *y* corresponds to regions to the right of the focal individual, whereas negative *y* corresponds to regions to the left of the focal individual.

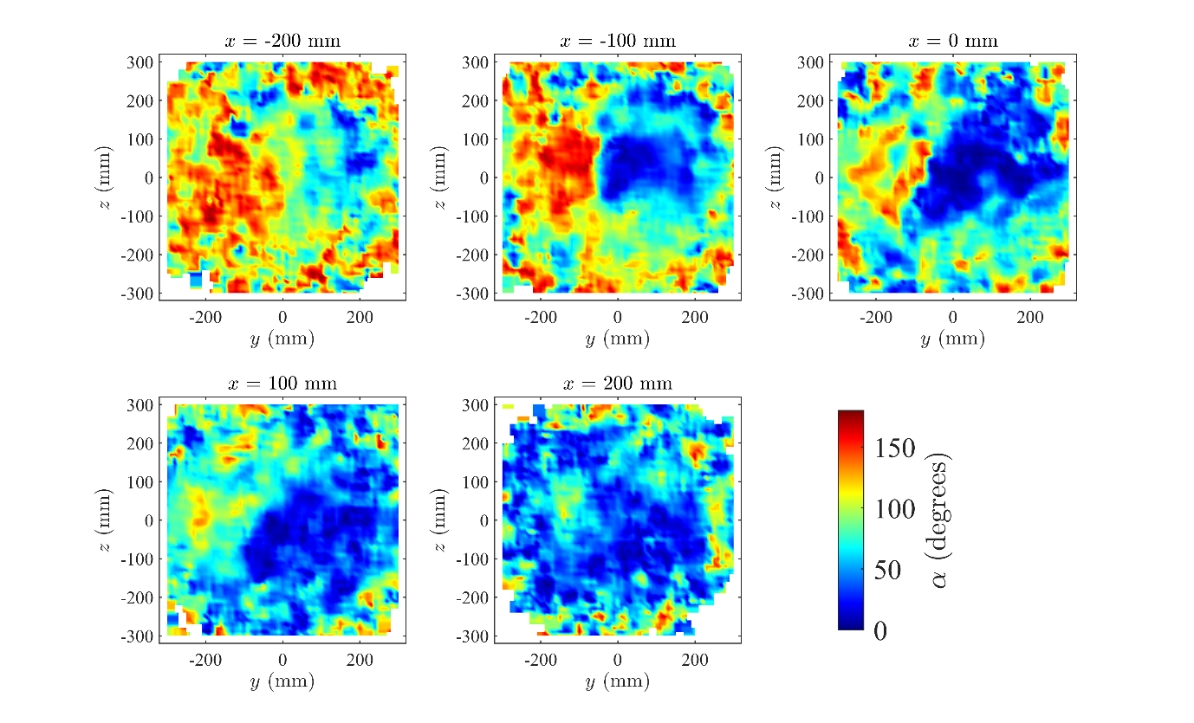

**FIGURE S11:** The mean angular difference in travelling direction of neighboring krill as a function of their (*x*, *y*, *z*) coordinates relative to a focal krill at (0, 0, 0), in the planes where *x* = -200 mm, *x* = -100 mm, *x* = 0 mm, *x* = 100 mm, and *x* = 200 mm. Positive *x* corresponds to regions in front of the focal individual, whereas negative *x* corresponds to regions behind the focal individual.

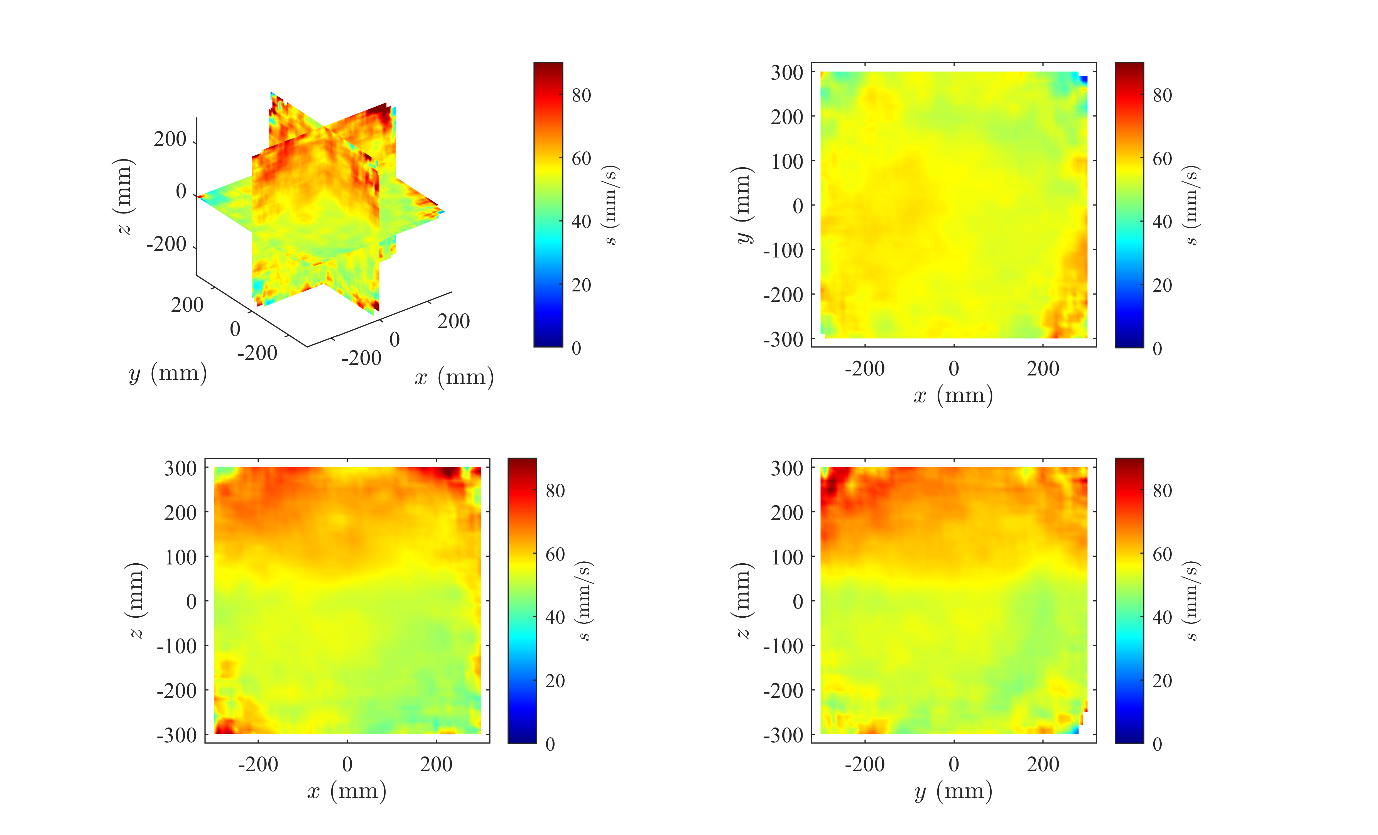

**FIGURE S12:** The mean speed of krill as a function of the relative (*x, y, z*) coordinates of neighbors, rendered in the planes *x* = 0, *y* = 0, and *z* = 0 (top left panel), and projected onto the *xy*- (top right panel), *xz*- (bottom left panel), and *yz*- (bottom right panel) planes. The colour scale for these plots has been truncated at 90 mm/s, such that the darkest red regions represent mean speeds greater than or equal to 90 mm/s.

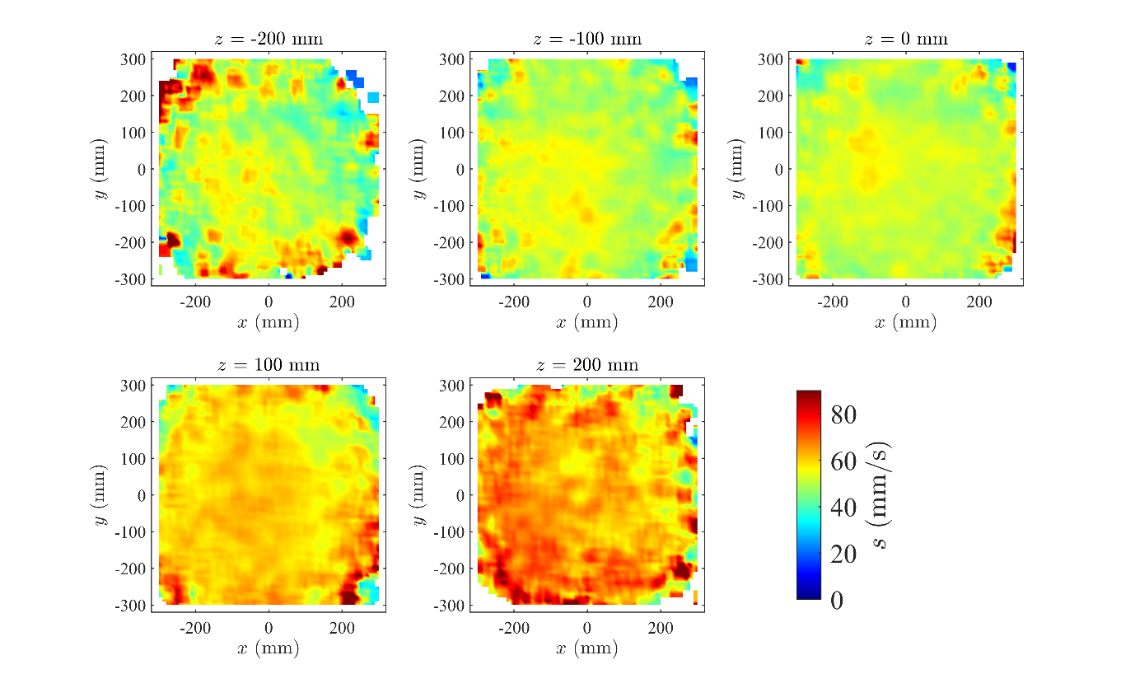

**FIGURE S13:** The mean speed of krill as a function of the relative (*x*, *y*, *z*) coordinates of their neighbors, in the planes where *z* = -200 mm, *z* = -100 mm, *z* = 0 mm, *z* = 100 mm, and *z* = 200 mm. The colour scale for these plots has been truncated at 90 mm/s, such that the darkest red regions represent mean speeds greater than or equal to 90 mm/s.

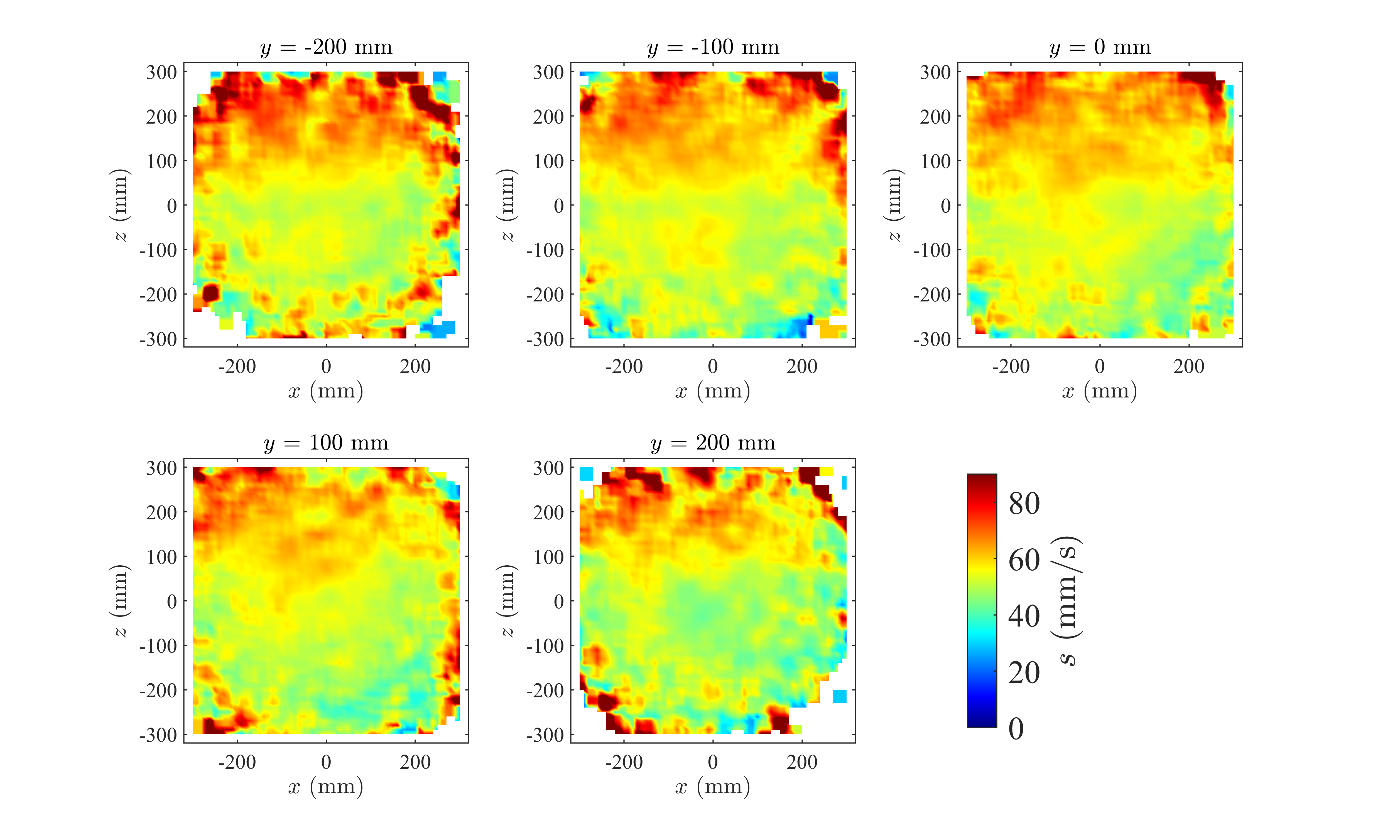

**FIGURE S14:** The mean speed of krill as a function of the relative (*x*, *y*, *z*) coordinates of their neighbors, in the planes where *y* = -200 mm, *y* = -100 mm, *y* = 0 mm, *y* = 100 mm, and *y* = 200 mm. The colour scale for these plots has been truncated at 90 mm/s, such that the darkest red regions represent mean speeds greater than or equal to 90 mm/s.

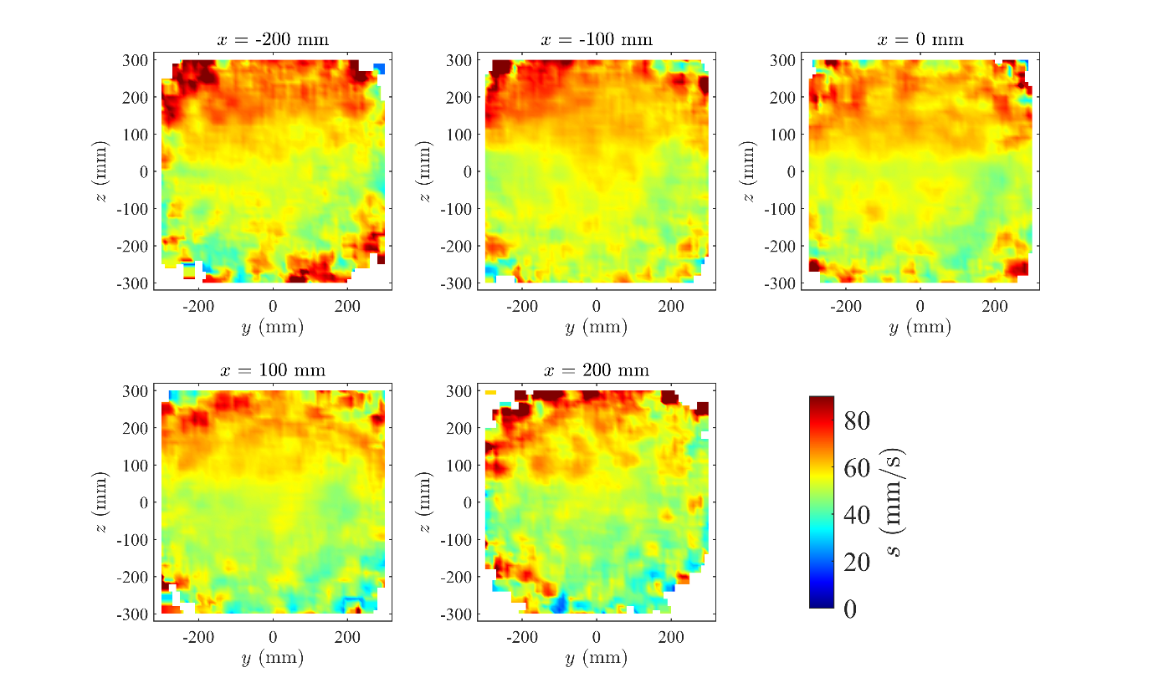

**FIGURE S15:** The mean speed of krill as a function of the relative (*x*, *y*, *z*) coordinates of their neighbors, in the planes where *x* = -200 mm, *x* = -100 mm, *x* = 0 mm, *x* = 100 mm, and *x* = 200 mm. The colour scale for these plots has been truncated at 90 mm/s, such that the darkest red regions represent mean speeds greater than or equal to 90 mm/s.

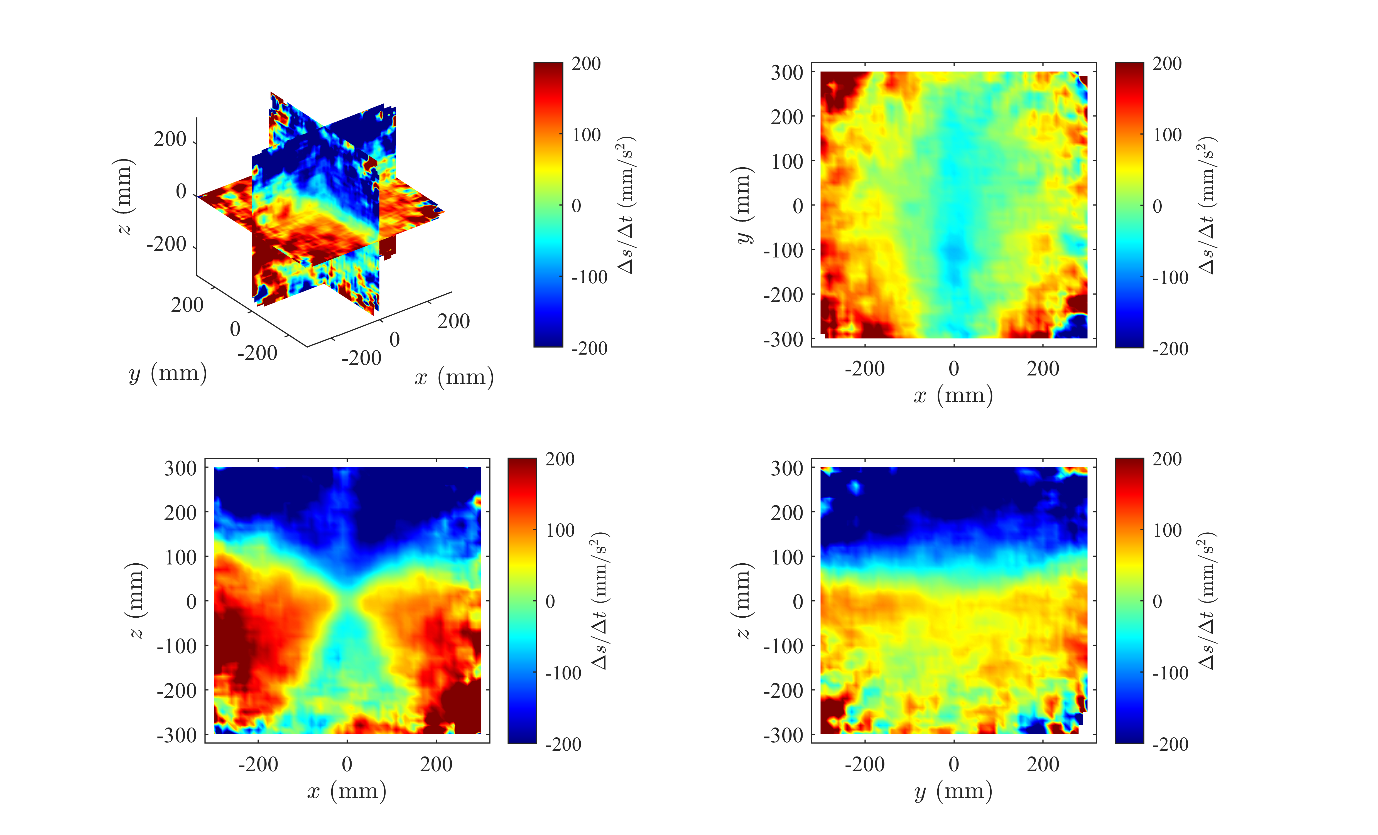

**FIGURE S16:** The mean change in speed over time of krill as a function of the relative (*x, y, z*) coordinates of neighbors, rendered in the planes *x* = 0, *y* = 0, and *z* = 0 (top left panel), and projected onto the *xy*- (top right panel), *xz*- (bottom left panel), and *yz*- (bottom right panel) planes. The colour scale for these plots has been truncated at ± 200 mm/s^2^, such that the darkest red regions represent mean increases in speed greater than or equal to 200 mm/s^2^, and the darkest blue regions represent mean decreases in speed less than or equal to 200 mm/s^2^.

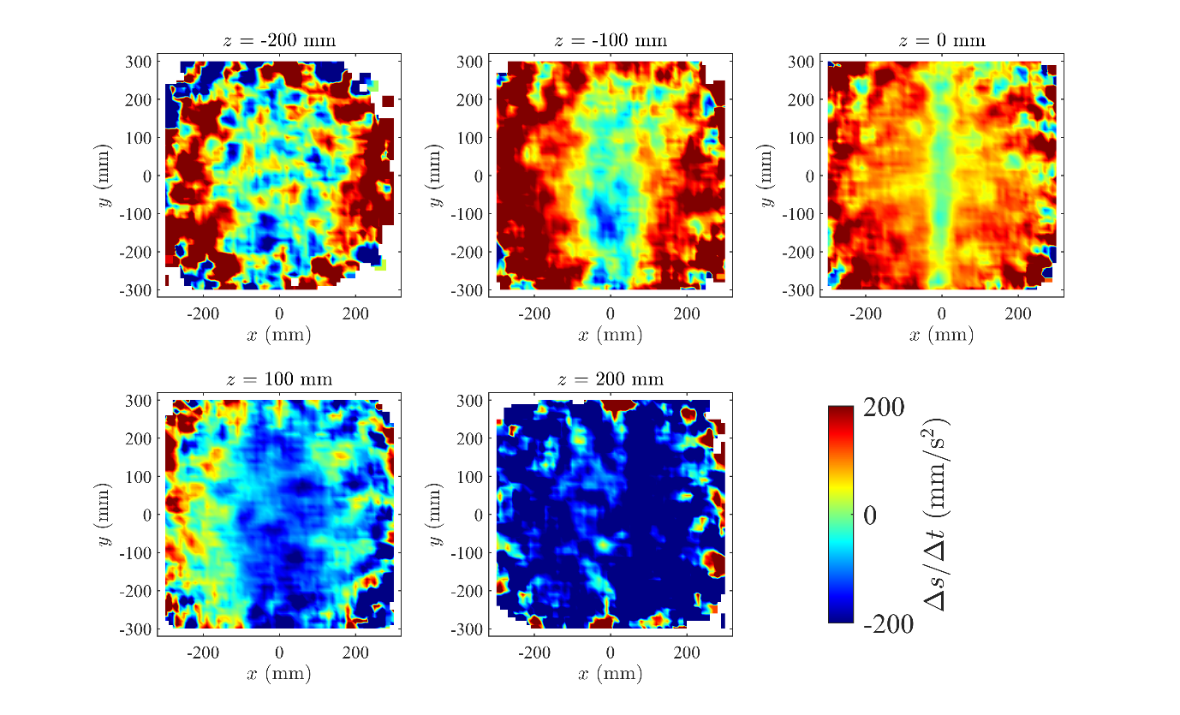

**FIGURE S17:** The mean change in speed over time of krill as a function of the relative (*x*, *y*, *z*) coordinates of their neighbors, in the planes where *z* = -200 mm, *z* = -100 mm, *z* = 0 mm, *z* = 100 mm, and *z* = 200 mm. The colour scale for these plots has been truncated at ± 200 mm/s^2^, such that the darkest red regions represent mean increases in speed greater than or equal to 200 mm/s^2^, and the darkest blue regions represent mean decreases in speed less than or equal to 200 mm/s^2^.

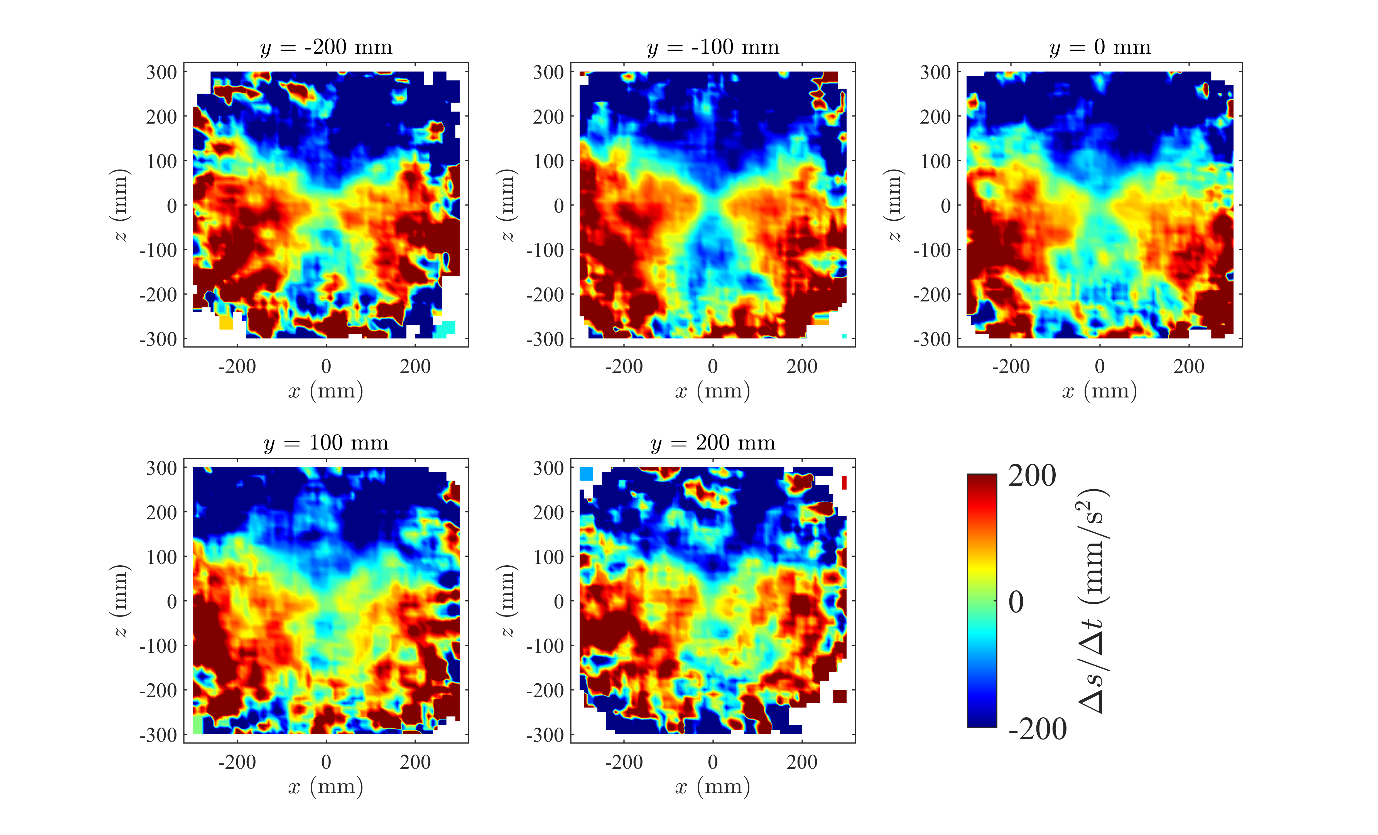

**FIGURE S18:** The mean change in speed over time of krill as a function of the relative (*x*, *y*, *z*) coordinates of their neighbors, in the planes where *y* = -200 mm, *y* = -100 mm, *y* = 0 mm, *y* = 100 mm, and *y* = 200 mm. The colour scale for these plots has been truncated at ± 200 mm/s^2^, such that the darkest red regions represent mean increases in speed greater than or equal to 200 mm/s^2^, and the darkest blue regions represent mean decreases in speed less than or equal to 200 mm/s^2^.

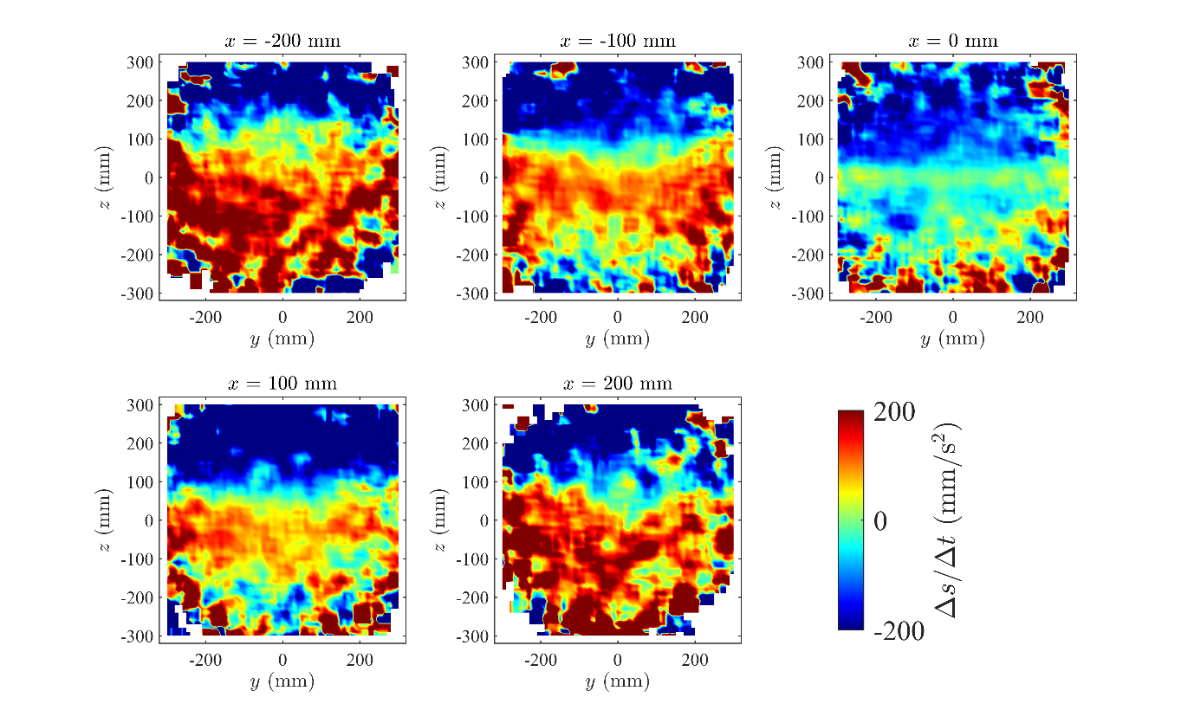

**FIGURE S19:** The mean change in speed over time of krill as a function of the relative (*x*, *y*, *z*) coordinates of their neighbors, in the planes where *x* = -200 mm, *x* = -100 mm, *x* = 0 mm, *x* = 100 mm, and *x* = 200 mm. The colour scale for these plots has been truncated at ± 200 mm/s^2^, such that the darkest red regions represent mean increases in speed greater than or equal to 200 mm/s^2^, and the darkest blue regions represent mean decreases in speed less than or equal to 200 mm/s^2^.

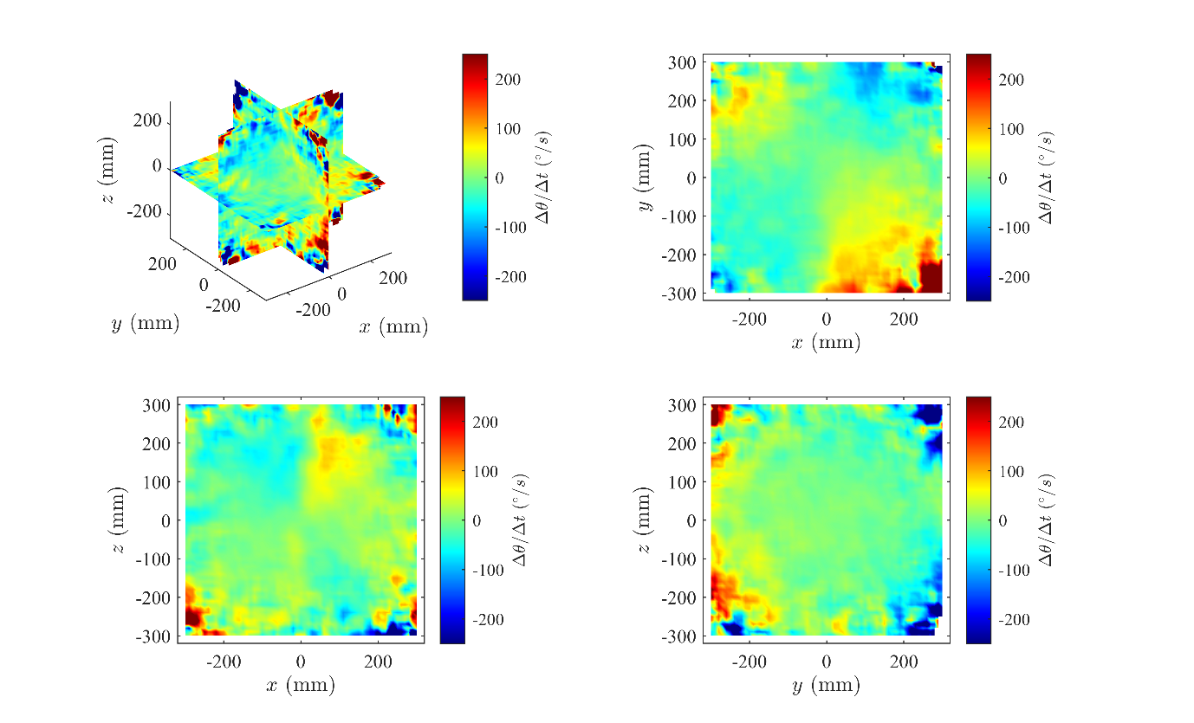

**FIGURE S20:** The component of turning in the plane *z* = 0, Δθ/Δt, of individuals as a function of the relative (*x*, *y*, *z*) coordinates of neighboring krill, rendered in the planes *x* = 0, *y* = 0, and *z* = 0 (top left panel), and projected onto the *xy-* (top right panel), *xz-* (bottom left panel), and *yz-*planes. The colour scale for these plots has been truncated at ± 250 degrees per second, such that the darkest red regions represent mean turns with leftward components at rates greater than or equal to 250 degrees per second, and the darkest blue regions represent turns with rightward components at rates greater than or equal to 250 degrees per second.

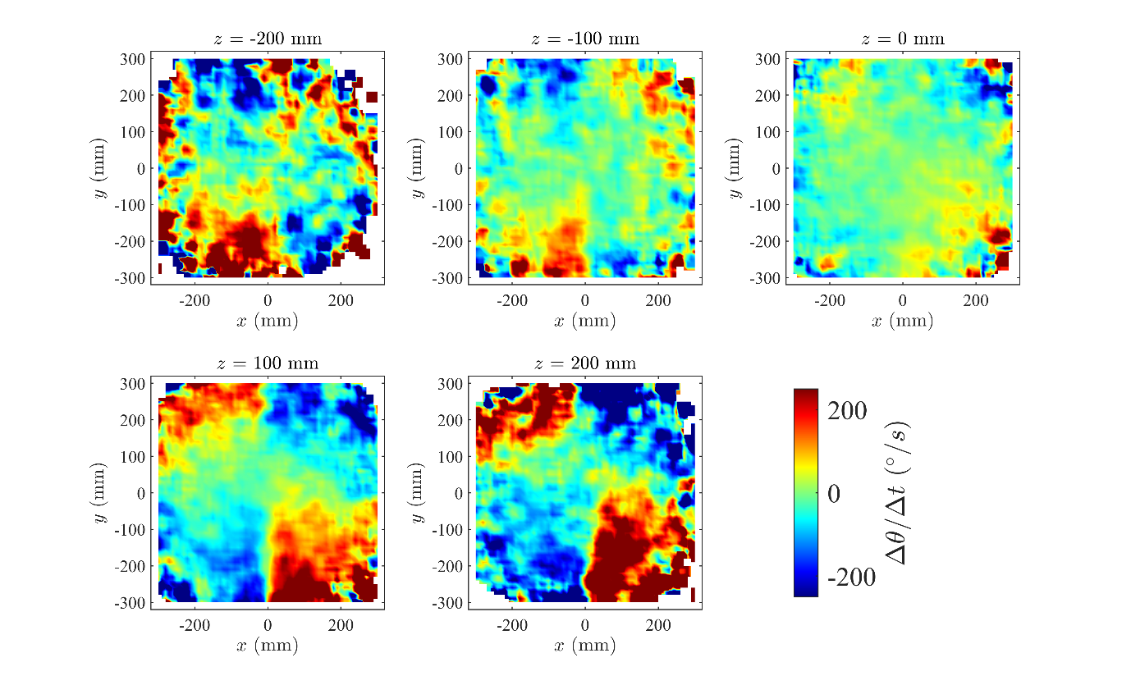

**FIGURE S21:** The component of turning in the plane *z* = 0, Δθ/Δt, of individuals as a function of the relative (*x*, *y*, *z*) coordinates of neighboring krill, in the planes where *z* = -200 mm, *z* = -100 mm, *z* = 0 mm, *z* = 100 mm, and *z* = 200 mm. The colour scale for these plots has been truncated at ± 250 degrees per second, such that the darkest red regions represent mean turns with leftward components at rates greater than or equal to 250 degrees per second, and the darkest blue regions represent turns with rightward components at rates greater than or equal to 250 degrees per second.

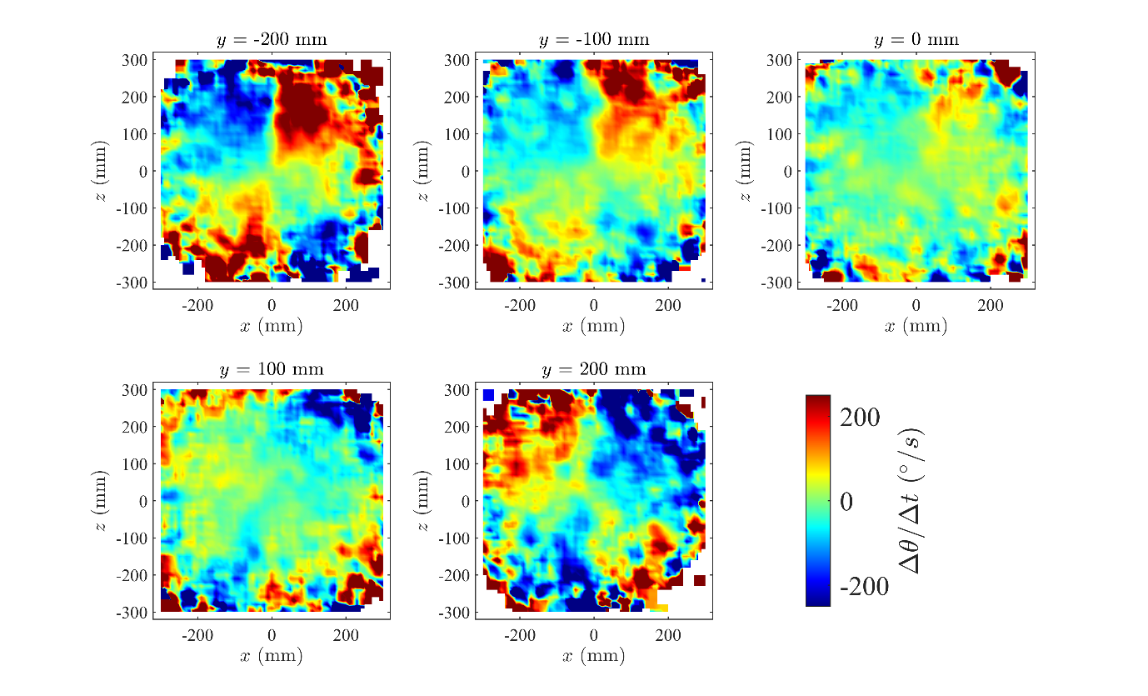

**FIGURE S22:** The component of turning in the plane *z* = 0, Δθ/Δt, of individuals as a function of the relative (*x*, *y*, *z*) coordinates of neighboring krill, in the planes where *y* = -200 mm, *y* = -100 mm, *y* = 0 mm, *y* = 100 mm, and *y* = 200 mm. The colour scale for these plots has been truncated at ± 250 degrees per second, such that the darkest red regions represent mean turns with leftward components at rates greater than or equal to 250 degrees per second, and the darkest blue regions represent turns with rightward components at rates greater than or equal to 250 degrees per second.

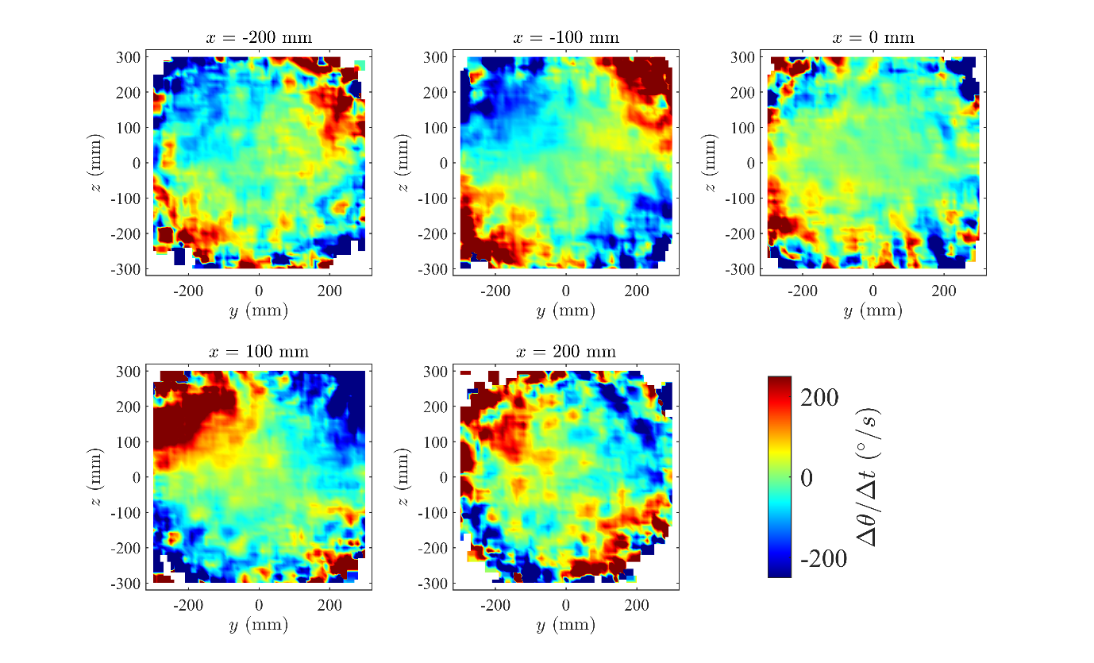

**FIGURE S23:** The component of turning in the plane *z* = 0, Δθ/Δt, of individuals as a function of the relative (*x*, *y*, *z*) coordinates of neighboring krill, in the planes where *x* = -200 mm, *x* = -100 mm, *x* = 0 mm, *x* = 100 mm, and *x* = 200 mm. The colour scale for these plots has been truncated at ± 250 degrees per second, such that the darkest red regions represent mean turns with leftward components at rates greater than or equal to 250 degrees per second, and the darkest blue regions represent turns with rightward components at rates greater than or equal to 250 degrees per second.

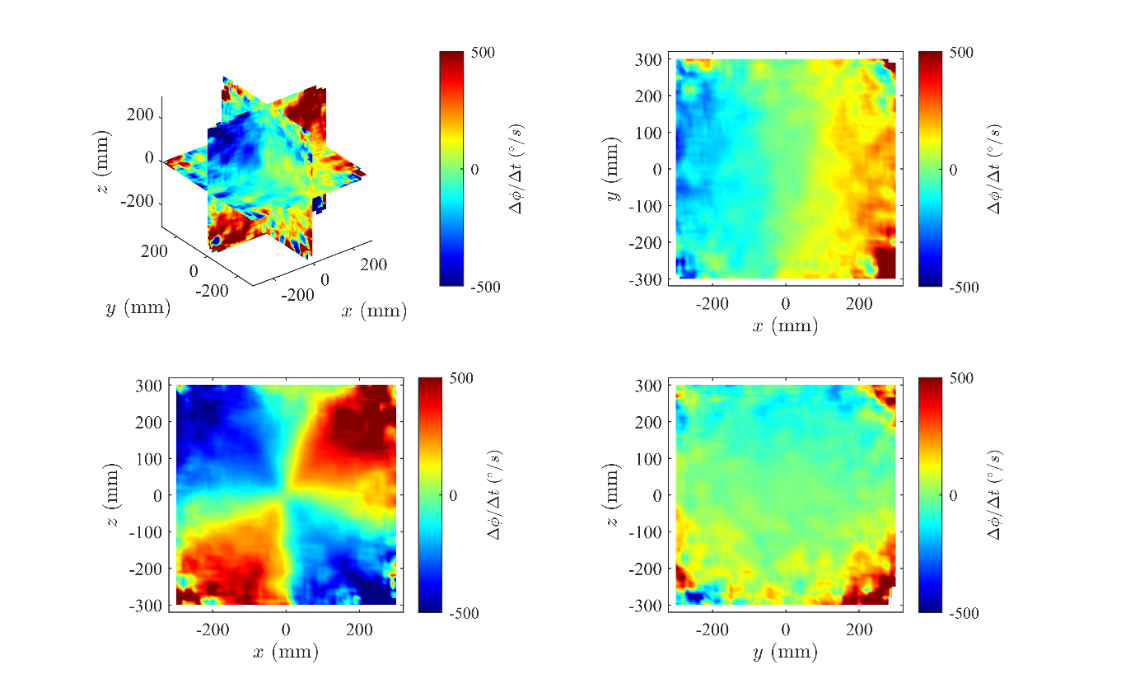

**FIGURE S24:** The component of turning perpendicular to the plane *z* = 0, Δ*ϕ*/Δt, of individuals as a function of the relative (*x*, *y*, *z*) coordinates of neighboring krill, rendered in the planes *x* = 0, *y* = 0, and *z* = 0 (top left panel), and projected onto the *xy-* (top right panel), *xz-* (bottom left panel), and *yz-*planes. The colour scale for these plots has been truncated at ± 500 degrees per second, such that the darkest red regions represent mean turns with upward components at rates greater than or equal to 500 degrees per second, and the darkest blue regions represent turns with downward components at rates greater than or equal to 500 degrees per second.

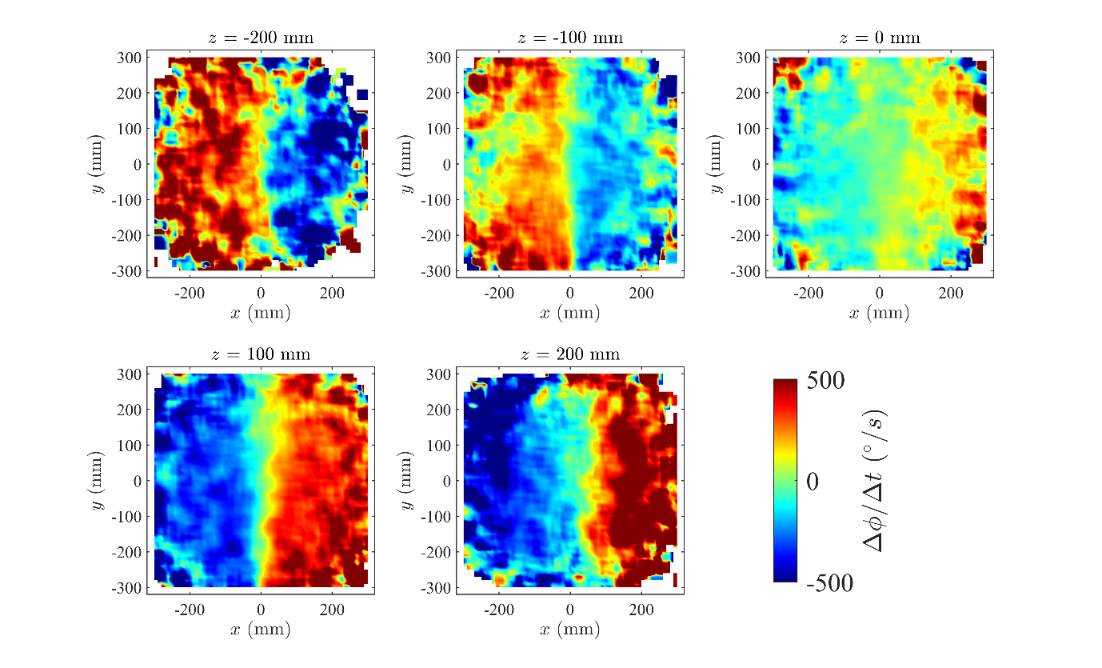

**FIGURE S25:** The component of turning perpendicular to the plane *z* = 0, Δ*ϕ*/Δt, of individuals as a function of the relative (*x*, *y*, *z*) coordinates of neighboring krill, in the planes where *z* = -200 mm, *z* = -100 mm, *z* = 0 mm, *z* = 100 mm, and *z* = 200 mm. The colour scale for these plots has been truncated at ± 500 degrees per second, such that the darkest red regions represent mean turns with upward components at rates greater than or equal to 500 degrees per second, and the darkest blue regions represent turns with downward components at rates greater than or equal to 500 degrees per second.

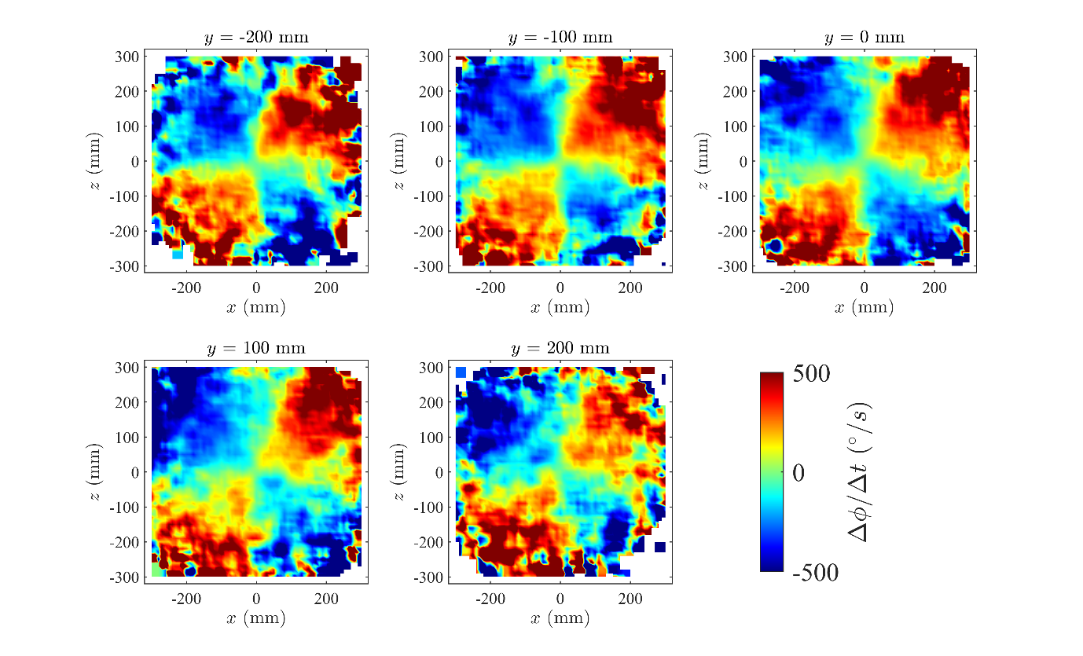

**FIGURE S26:** The component of turning perpendicular to the plane *z* = 0, Δ*ϕ*/Δt, of individuals as a function of the relative (*x*, *y*, *z*) coordinates of neighboring krill, in the planes where *y* = -200 mm, *y* = -100 mm, *y* = 0 mm, *y* = 100 mm, and *y* = 200 mm. The colour scale for these plots has been truncated at ± 500 degrees per second, such that the darkest red regions represent mean turns with upward components at rates greater than or equal to 500 degrees per second, and the darkest blue regions represent turns with downward components at rates greater than or equal to 500 degrees per second.

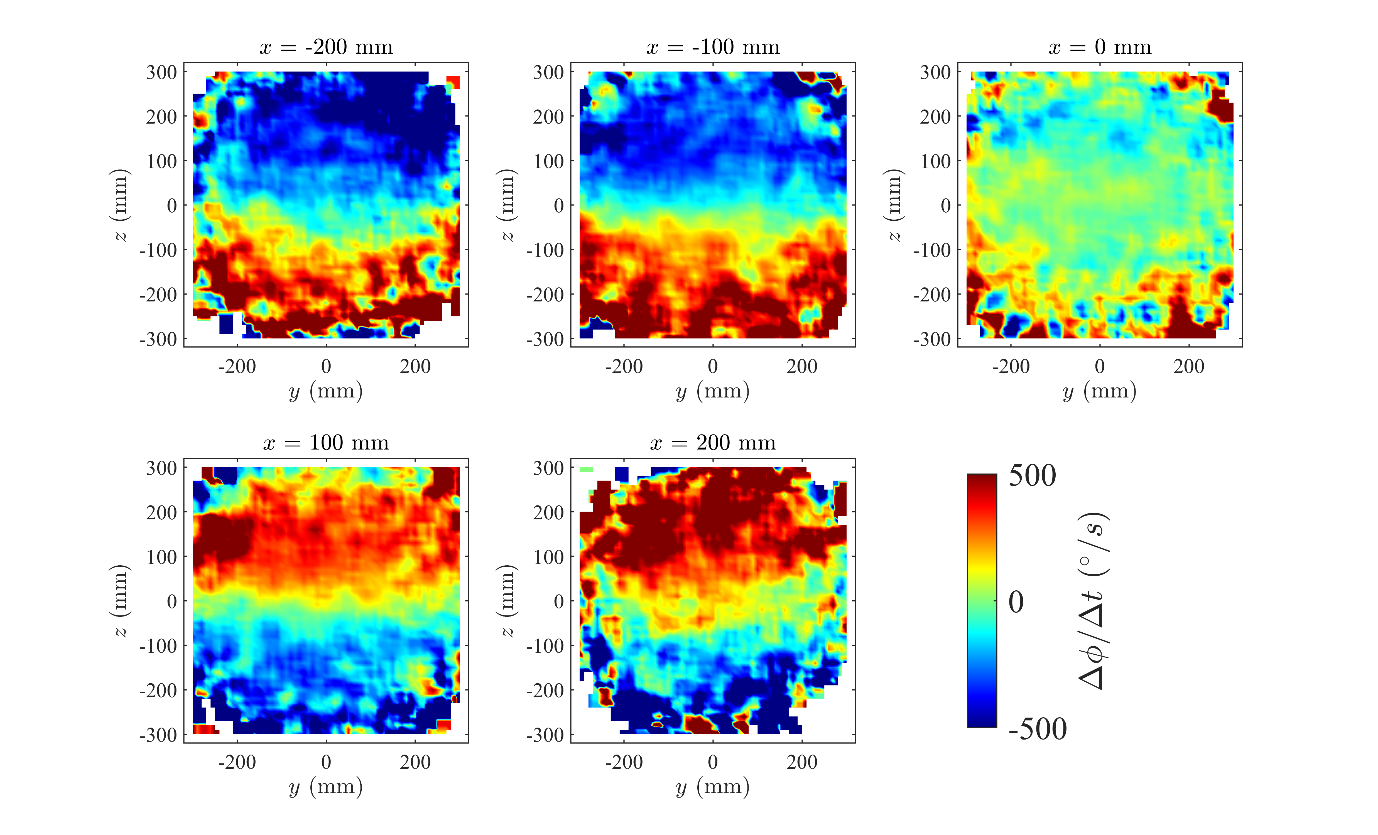

**FIGURE S27:** The component of turning perpendicular to the plane *z* = 0, Δ*ϕ*/Δt, of individuals as a function of the relative (*x*, *y*, *z*) coordinates of neighboring krill, in the planes where *x* = -200 mm, *x* = -100 mm, *x* = 0 mm, *x* = 100 mm, and *x* = 200 mm. The colour scale for these plots has been truncated at ± 500 degrees per second, such that the darkest red regions represent mean turns with upward components at rates greater than or equal to 500 degrees per second, and the darkest blue regions represent turns with downward components at rates greater than or equal to 500 degrees per second.

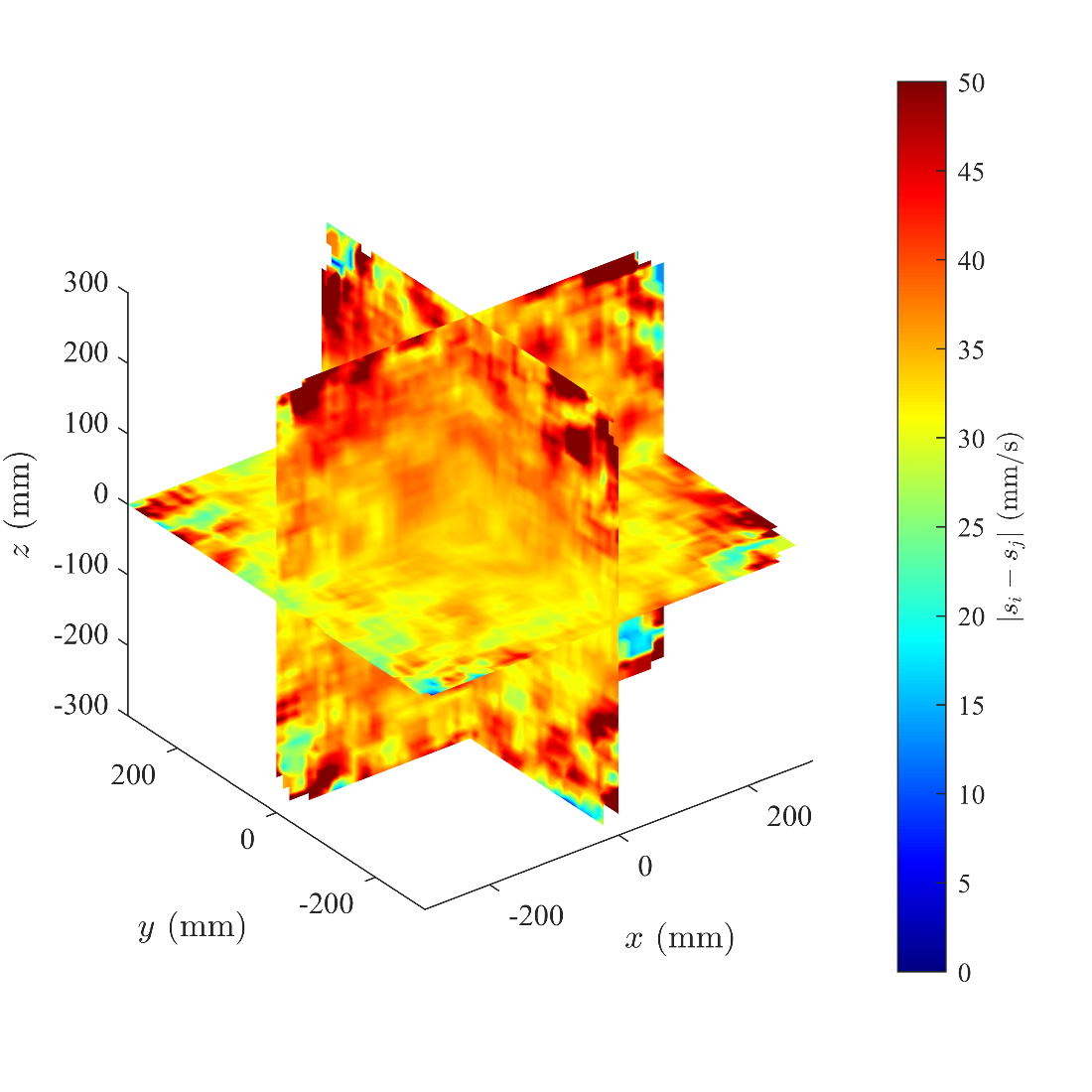

**FIGURE S28:** The mean absolute difference in speed between individuals *i* located at the origin and their neighbors *j* located at coordinate (*x*, *y*, *z*), rendered in the planes *x* = 0, *y* = 0, and *z* = 0. The colour scale is truncated at 50 mm/s, so regions of the darkest red colour indicate speed differences greater than or equal to 50 mm/s.

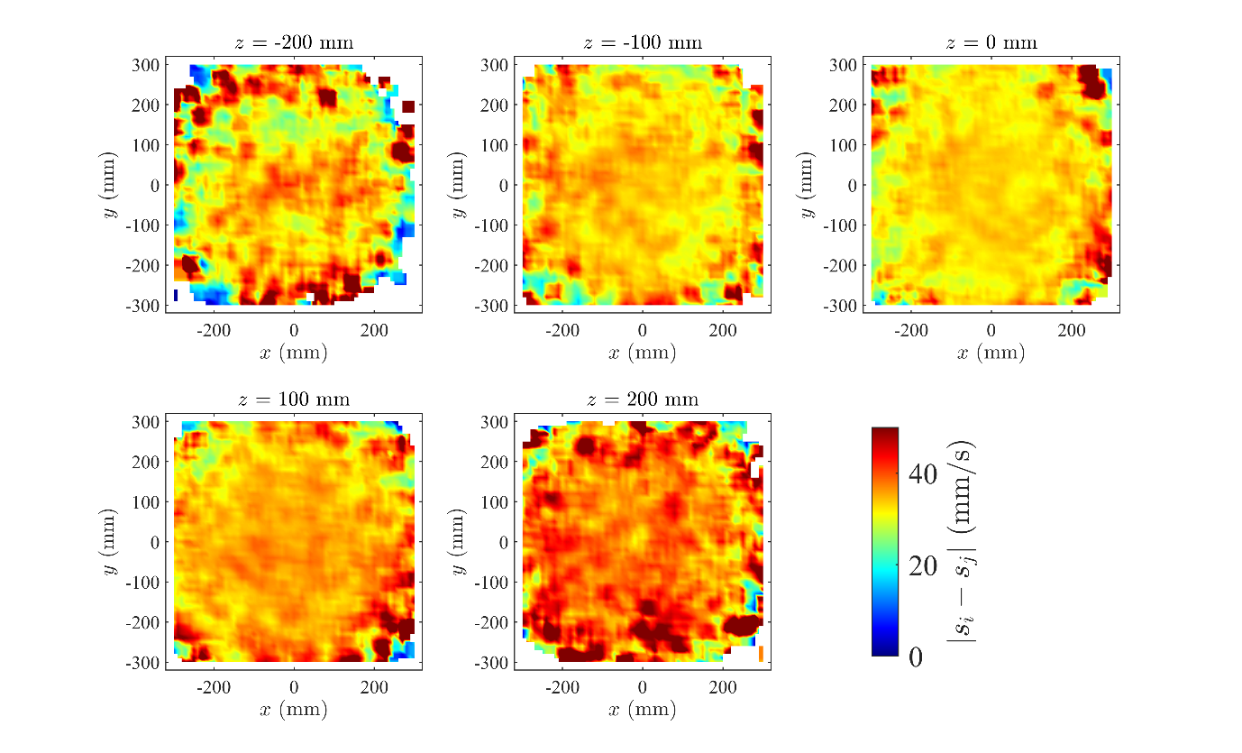

**FIGURE S29:** The mean absolute difference in speed between individuals *i* located at the origin and their neighbors *j* located at coordinate (*x*, *y*, *z*), rendered in the planes where *z* = -200 mm, *z* = -100 mm, *z* = 0 mm, *z* = 100 mm, and *z* = 200 mm. The colour scale is truncated at 50 mm/s, so regions of the darkest red colour indicate speed differences greater than or equal to 50 mm/s.

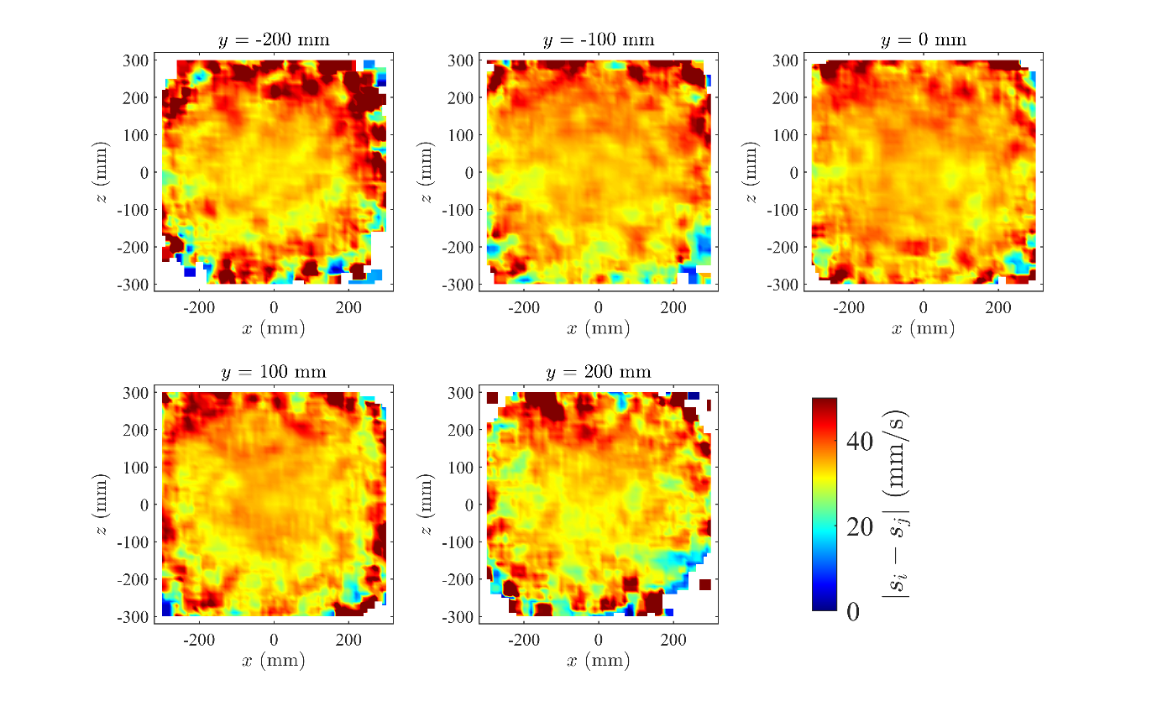

**FIGURE S30:** The mean absolute difference in speed between individuals *i* located at the origin and their neighbors *j* located at coordinate (*x*, *y*, *z*), rendered in the planes where *y* = -200 mm, *y* = -100 mm, *y* = 0 mm, *y* = 100 mm, and *y* = 200 mm. The colour scale is truncated at 50 mm/s, so regions of the darkest red colour indicate speed differences greater than or equal to 50 mm/s.

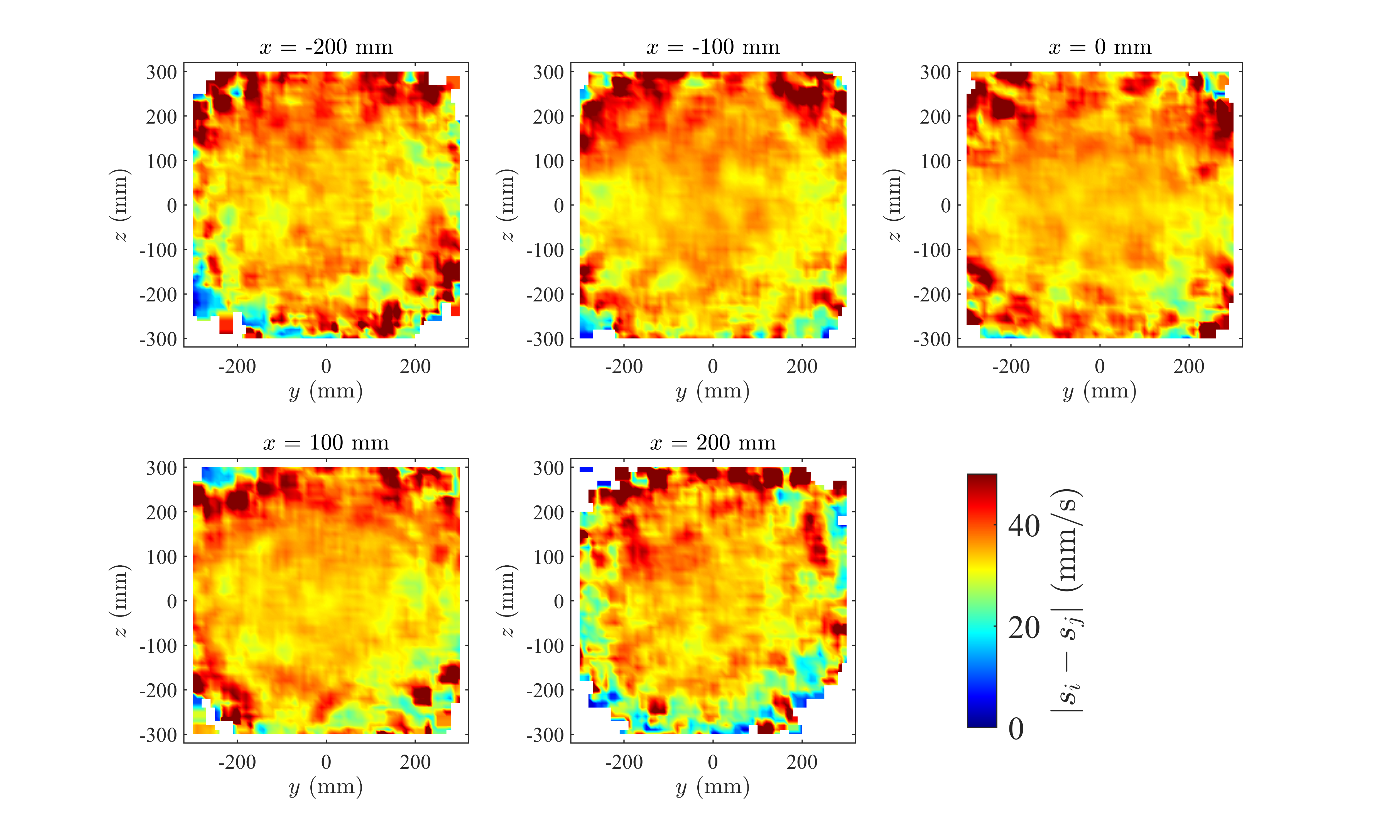

**FIGURE S31:** The mean absolute difference in speed between individuals *i* located at the origin and their neighbors *j* located at coordinate (*x*, *y*, *z*), rendered in the planes where *x* = -200 mm, *x* = -100 mm, *x* = 0 mm, *x* = 100 mm, and *x* = 200 mm. The colour scale is truncated at 50 mm/s, so regions of the darkest red colour indicate speed differences greater than or equal to 50 mm/s.

**Table S1**: Optimized parameters for TE calculations

| Parameter | Samples | k | tau | lag | mean TE | surrogates | null distribution of surrogates | p(surrogate > measured) |
| --- | --- | --- | --- | --- | --- | --- | --- | --- |
| Speed | 515639 | 5 | 1 | 3 | -0.001 | 100 | -0.0084 +/- 0.0007 | 0.00000 |
| Heading | 533766 | 2 | 1 | 1 | 0.062 | 100 | -0.0022 +/- 0.0008 | 0.00000 |

**Table S2:** Broad qualitative outline of the signs of the observed components of turning, Δθ/Δt and Δ*ϕ*/Δt, when neighboring krill occupied given octants of three dimensional space (derived from Figures S21 and S25).

| *x* | *y* | *z* | Δθ/Δt | Δ*ϕ*/Δt |
| --- | --- | --- | --- | --- |
| > 0 | > 0 | > 0 | < 0 (rightward, away from neighbor) | > 0 (upward, towards neighbor) |
| < 0 | > 0 | > 0 | > 0 (leftward, towards neighbor) | < 0 (downward, away from neighbor) |
| > 0 | < 0 | > 0 | > 0 (leftward, away from neighbor) | > 0 (upward, towards neighbor) |
| < 0 | < 0 | > 0 | < 0 (rightward, towards neighbor) | < 0 (downward, away from neighbor) |
| > 0 | > 0 | < 0 | > 0 (leftward, towards neighbor) | < 0 (downward, towards neighbor) |
| < 0 | > 0 | < 0 | < 0 (rightward, away from neighbor) | > 0 (upward, away from neighbor) |
| > 0 | < 0 | < 0 | < 0 (rightward, towards neighbor) | < 0 (downward, towards neighbor) |
| < 0 | < 0 | < 0 | > 0 (leftward, away from neighbor) | > 0 (upward, away from neighbor) |
